# A lipid fenestration-gated membrane depolarization mechanism expands the repertoire of CRISPR-mediated anti-phage defense strategies

**DOI:** 10.64898/2026.08.05.743128

**Authors:** Puja Majumder, Heloise Carion, Gianna Stella, Dinshaw J Patel, Luciano A Marraffini

**Author notes:** These authors contributed equally (Puja Majumder, Heloise Carion).

## Abstract

Prokaryotic Type III CRISPR–Cas systems synthesize cyclic oligoadenylate (cOA) second messengers that activate CRISPR-associated Rossmann fold (CARF) immune effectors during viral infection. Here we characterize Chp1, a membrane-embedded CARF effector that assembles as a tetrameric pore harboring autoinhibitory lipid fenestrations that occlude the pore in the inactive state. cOA binding triggers a structural rearrangement that (i) closes the fenestrations, (ii) eliminates the lipid obstruction of the pore to open it, and (iii) remodels the pore entrance from hydrophobic to polar to allow ion permeation. This conformational switch triggers depolarization of the membrane of the infected cell, which enters growth arrest and becomes inhospitable for viral replication. Our results uncover a lipid-gated mechanism that expands the repertoire of CRISPR-mediated defense strategies.

## INTRODUCTION

Cell membranes constitute a barrier between the cytosol and the extracellular environment, enforcing stringent control over the intracellular ionic composition (*1*, *2*). The permeability of specific solutes requires specialized membrane proteins that function as channels and transporters (*3*). Protons are fundamental permeant ions that generate an electrochemical gradient across membranes (*4*, *5*). The energy accumulated in this gradient is then used for a broad range of core cellular processes across all domains of life, including ATP synthesis, nutrient uptake and motility (*6*, *7*). Bacteria exploit the essential nature of the membrane potential as a sophisticated antiphage defense strategy, inducing membrane perturbations or depolarization to disrupt this potential and cause a growth arrest of the infected host that prevents viral replication (*8*–*12*).

Clustered, regularly interspaced, short palindromic repeats (CRISPR) loci and their associated genes (*cas*) provide RNA-guided adaptive immunity to prokaryotes, protecting them from phage (*13*) and plasmid (*14*) infection. Depending on the *cas* gene content, these defense systems can be classified into seven different types, I-VII (*15*). In the case of type III CRISPR-Cas systems, immunity is provided through activation of CRISPR-associated Rossmann fold (CARF) effectors (*16*), some of which are located in the membrane and cause membrane depolarization (*17*–*19*). Type III CRISPR-*cas* loci encode the Cas10 complex that uses a guide RNA to find complementary invader transcripts produced during infection to trigger an immune response (*16*). Target binding induces the activity of the Palm domain of Cas10, which uses ATP to synthesize cyclic oligoadenylates (cOA), primarily cyclic tetra- and hexa-adenylates (cA_4_ and cA_6_) (*20*, *21*).

cOA act as second messengers that bind to CARF domains and stimulate diverse effector activities that are toxic to the infected host and thus prevent the propagation of the invader (*16*). To date, four CARF effectors containing transmembrane (TM) domains have been studied, all of which mediate type III CRISPR-Cas anti-phage immunity. Cap1 (*19*) and Cam1 (*18*) cause membrane depolarization when activated by cA_4_, in the case of Cap1 through a rearrangement of the TM domain that opens a pore. Csx23 also harbors a TM domain that suffers a conformational change upon ligand binding (*17*); however how it affects the host membrane is not currently well understood. Chp1 contains not only CARF and TM regions, but also a HAD phosphatase domain (*22*). Our initial characterization of this effector focused on the study of the CARF and HAD domains in isolation due to the challenges of purifying full-length Chp1. It was shown that the purified CARF domain binds cA_4_, and that the purified HAD domain dephosphorylates ATP, dATP and IMP to generate ADP, dADP and inosine, respectively (*22*). In vivo, activation of Chp1 in staphylococci results in the depletion of ATP and IMP upon activation of the type III-A CRISPR-Cas response.

Here we determined the cryo-EM structures of apo- and cA_4_-bound, full-length Chp1, in the presence or absence of the ATP analog ApCpp. Contrary to expectations from our previous analysis of the HAD domain, we found that ligand binding does not significantly alter the conformation of this domain, including the architecture of the ATP binding pocket. Biochemical assays showed that dephosphorylation of ATP and dATP by purified full-length Chp1 is similar to that of the isolated HAD domain and does not change in the presence of cA_4_. Instead, ligand binding causes the rearrangement of a linker helix coupled to the TM region. This region forms a pore whose center is occluded by membrane lipids that enter through lateral tunnels, known as fenestrations, in the inactive state. In the active state, the switch of the linker helix leads to the closing of the fenestrations and the exclusion of the membrane lipids from the pore center, thus opening the pore. In addition, the environment of the cytosolic entry of the pore changes from hydrophobic to polar, presumably to enable ion conduction. Consistent with these new results, Chp1 activation causes membrane depolarization in vivo. Collectively, our results have uncovered a distinct and unanticipated mechanism of pore activation in which second messenger signaling of viral infection enables allosteric control of pore opening via regulation of lipid fenestrations.

## RESULTS

### Chp1 forms a transmembrane tetrameric complex

*Bacteroidales* bacterium Chp1 exhibits a tripartite architecture comprising an N-terminal HAD domain, a central four-helix transmembrane domain (TM1-4), and a C-terminal CARF domain (**Fig. 1A**). To characterize the function of Chp1 at the atomic level, we decided to obtain the structure of this CARF effector. We added an N-terminal His₆-tag to Chp1, expressed in *E. coli* and purified it in the presence of glyco-diosgenin (GDN) detergent to increase the solubility of the transmembrane regions of the protein (**Figs. S1A, B**). Size-exclusion chromatography coupled with multi-angle light scattering (SEC–MALS) indicated that Chp1 assembles as a homotetramer in solution (**Fig. S1C**). Single-particle cryo-electron microscopy (cryo-EM) resolved two apo-Chp1 classes (class A, **Figs. 1B, S1D**, and B, **Fig. S1E**) that exhibit minimal conformational differences (**Fig. S1F**), indicating limited intrinsic flexibility in the absence of ligand. Both reconstructions revealed a tetrameric architecture in which the TM helices are embedded within the GDN detergent micelle (**Figs. 1B, S1D,E**). Given the previous finding that Chp1 localizes to the staphylococcal membrane fraction (*22*), we conclude that this domain is inserted into the bacterial lipid bilayer. The individual structure of subunit 1 (**Fig. S1G**) shows the cytosolic N-terminal HAD domain connected to TM1 via the flexible linker L1. TM1 spans the membrane towards the extracellular side and is followed by TM2, which bends back towards the cytosol. TM2 is linked to TM3 through a 22-residue loop-helix-loop segment (segment J) that lies along the cytosolic membrane surface but does not insert into the membrane. TM3 traverses the membrane to the extracellular side and is followed by TM4, which returns towards the cytosol and connects to the C-terminal CARF domain via a second linker (L2) (**Fig. S1G**).

**Figure 1.**
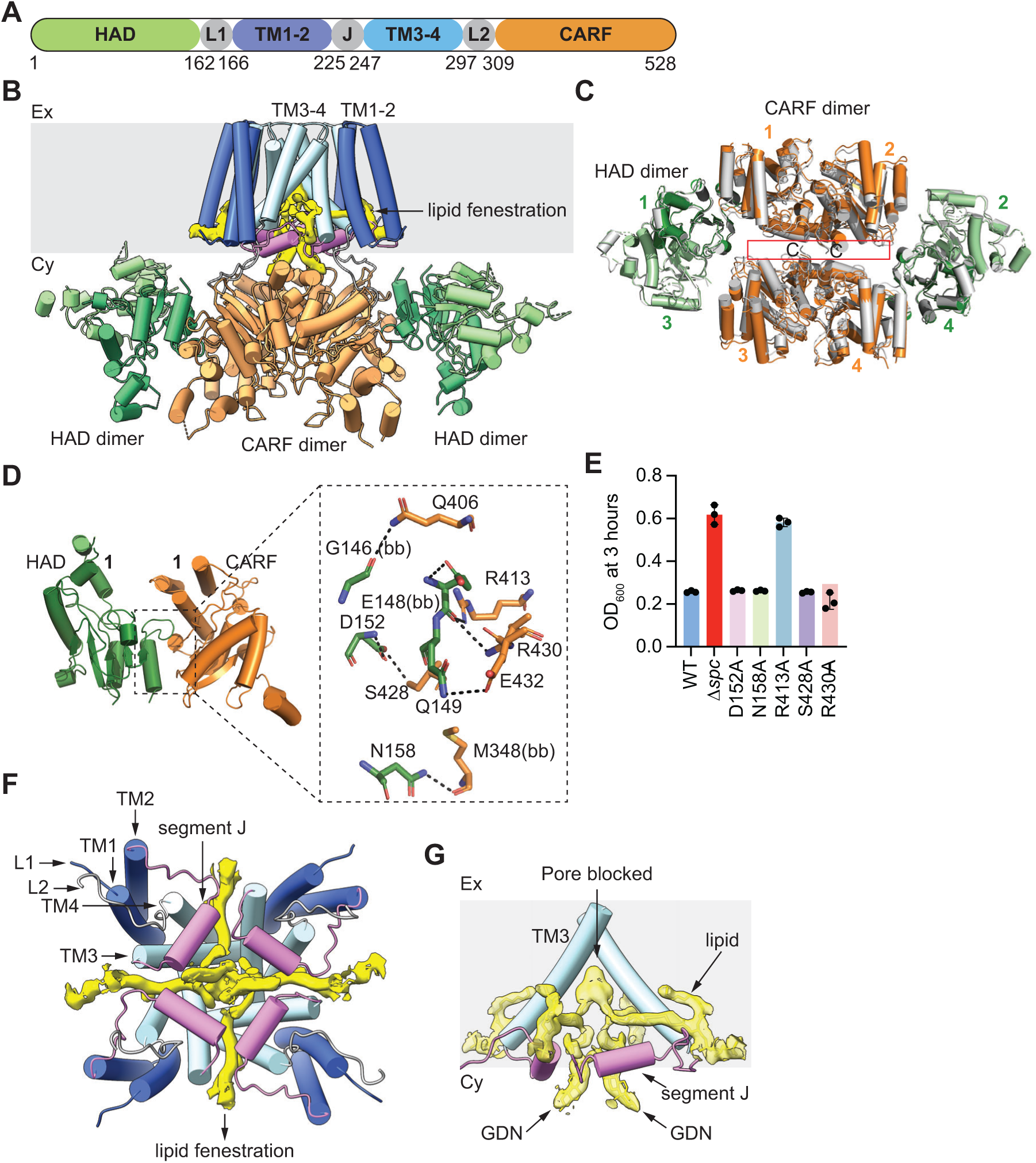
Inactive Chp1 forms a tetrameric transmembrane pore closed by lipid fenestrations. **(A)** Full-length Chp1 comprises an N-terminal HAD domain (green) linked to transmembrane helices TM1–2 (blue) via linker 1 (L1, grey). TM1–2 connects to TM3–4 (light blue) through segment J (violet), and TM3–4 is joined to the C-terminal CARF domain (orange) by linker 2 (L2, grey). **(B)** Cryo-EM map (class A) of apo Chp1 displaying a tetrameric organization, with both the HAD and CARF domains located in the cytosol, while the TM domain anchors the protein to the membrane region (grey area). Lipid fenestrations are shown in yellow in the TM domain. **(C)** Superposition of the cytosolic domain of apo Chp1 class A (color) and class B (silver). The red box shows the CARF tetrameric interface formed by a pair of CARF dimers. Numbers indicate the CARF (orange) and HAD (green) domains of each of the four Chp1 subunits. **(D)** HAD-CARF domains of subunit 1 from apo Chp1 class A structure displaying the interdomain interacting residues. (bb) indicates backbone interaction. **(E)** Growth of staphylococci carrying pTarget and pCRISPR variants harboring alanine substitutions of Chp1 residues shown in **(D)**, measured as the OD_600_ value 3 hours after addition of aTc. Data are mean of three biological replicates ± s.e.m. **(F)** Cytosolic view of the TM domain highlighting the arrangement of TM1–TM4, segment J, L1 and L2. A central pore lined by TM3 and segment J is occupied by lipid fenestrations (yellow). **(G)** Side view of the TM domain highlighting the lipid fenestration entering and blocking the central pore of Chp1 at the resting state. Lipid and GDN detergent maps are shown in yellow.

On the cytosolic side, the CARF domains of subunits 1 and 2, and 3 and 4, form dimers that assemble into a central dimer-of-dimers topology, flanked by two HAD dimers one formed by subunits 1 and 3 and the other by subunits 2 and 4, positioned on opposing sides of the complex, thereby cross-bracing the tetrameric assembly (**Figs. 1C, S2A,B**). Differential interactions between these domains establish asymmetry within the complex. Whereas the CARF and HAD domains are spatially separated in subunits 2 and 3 (cyan arrowheads, **Figs. S2A,B**), there are direct contacts between the N-terminal and C-terminal regions of the HAD and CARF domains, respectively, in subunits 1 and 4 (red arrowheads, **Figs. S2A,B**). These interactions (**Fig. 1D**) should mediate the stabilization of the cytosolic assembly, since there are sparse contacts between CARF dimers (red rectangle, **Fig. 1C**).

To test the importance of these interfaces, we introduced alanine substitutions at the key residues involved and evaluated Chp1-mediated toxicity in vivo. We performed an assay that artificially activates CARF effectors in the absence of phage infection (*23*), using staphylococci carrying pCRISPR and pTarget plasmids. pCRISPR encodes the *Staphylococcus epidermidis* RP62a type III-A CRISPR-Cas system along with a CARF effector such as Chp1 (**Fig. S2C**). pTarget produces a target RNA whose transcription is regulated by an anhydrotetracycline (aTc) inducible promoter. Upon addition of aTc to the bacterial culture, recognition of the target RNA by the Cas10 complex results in the synthesis of cOA that trigger the toxic activity of CARF effectors (*20*, *21*). First, we used this assay to evaluate the importance of the dimerization of the CARF and HAD domains for Chp1 activity in vivo. As previously reported (*22*), Chp1 activation after aTc addition resulted in a marked growth arrest, not observed in the absence of a targeting spacer (Δ*spc*), determined by measuring the optical density at 600 nm (OD_600_) of the cultures over time. We substituted the residues that participate in CARF dimerization (E442, Y492, R506 and K525, **Fig. S2D**) for alanine; however none of the mutations affected Chp1-mediated toxicity in staphylococci (**Figs. S2E,F**). We obtained similarly negative results for alanine substitutions of residues that participate in interactions between HAD domains; H109A, S86A, I114A, L120A, C121A and E122A (**Fig. S2G**) had no detectable effects on Chp1 activity in vivo (**Figs. S2H,I**). We then evaluated the importance of the HAD-CARF interactions by testing the effect of alanine substitutions of the D152, N158, R413, S428 or R430 residues (**Fig. 1D**) and found that R413A led to a complete loss of Chp1-induced toxicity (**Figs. S3A, 1E**). These results highlight the importance of the asymmetric CARF-HAD inter-domain interactions in stabilizing the cytosolic region of the Chp1 tetramer.

### Lipid fenestrations occlude the central pore of inactive Chp1

The four TM helices of Chp1 are embedded within the GDN micelle (presumably also within the bacterial membrane, see above) and display interactions between residues of adjacent TM3 (W263, N266, L271, F272, E275, N276 and S283) and TM1 (H198) helices (**Fig. S3B**). Upon tetramer assembly, sixteen TM helices traverse the membrane (**Fig. 1B**) displaying four-fold symmetry, with TM3-TM4 forming a central core in which TM3 and segment J line a symmetric pore, and TM1–TM2 occupying peripheral positions at the four corners (**Fig. 1F**). Interestingly, cryo-EM revealed membrane lipid acyl chains penetrating the central pore through lateral openings in the transmembrane domain, consistent with the presence of lipid fenestrations (**Figs. 1B,F**). These are hydrophobic tunnels that enable the penetration of membrane lipids into the pore interior. Lipid fenestrations have been previously found in various membrane channels, including voltage-gated (sodium, potassium and calcium), mechanosensitive and two-pore potassium channels, where they provide lateral access pathways for lipid acyl chains and hydrophobic ligands into the pore cavity(*24*–*33*). The Chp1 cryo-EM structure showed four symmetric fenestrations formed by the TM3 helix and segment J that extend toward the central pore, through which lipid acyl chains and GDN detergent molecules penetrate to occupy the pore lumen, resulting in pore occlusion (**Figs. 1F,G**).

### Nucleotide binding at the HAD domain does not affect Chp1 topology

To further understand the ATPase activity of Chp1, we solved the cryo-EM structure of full-length apo-Chp1 bound to the non-hydrolysable ATP analog ApCpp, in which a carbon replaces the bridging oxygen at the α–β phosphodiester bond. We found one ApCpp molecule bound to each of the four HAD domains (**Fig. 2A**). Nucleotide binding, however, caused minimal rearrangement of the tetramer, and the alignment of the Chp1 in the absence and presence of ApCpp revealed close correspondence, with an RMSD of 1.4 Å (**Fig. 2B**).

**Figure 2.**
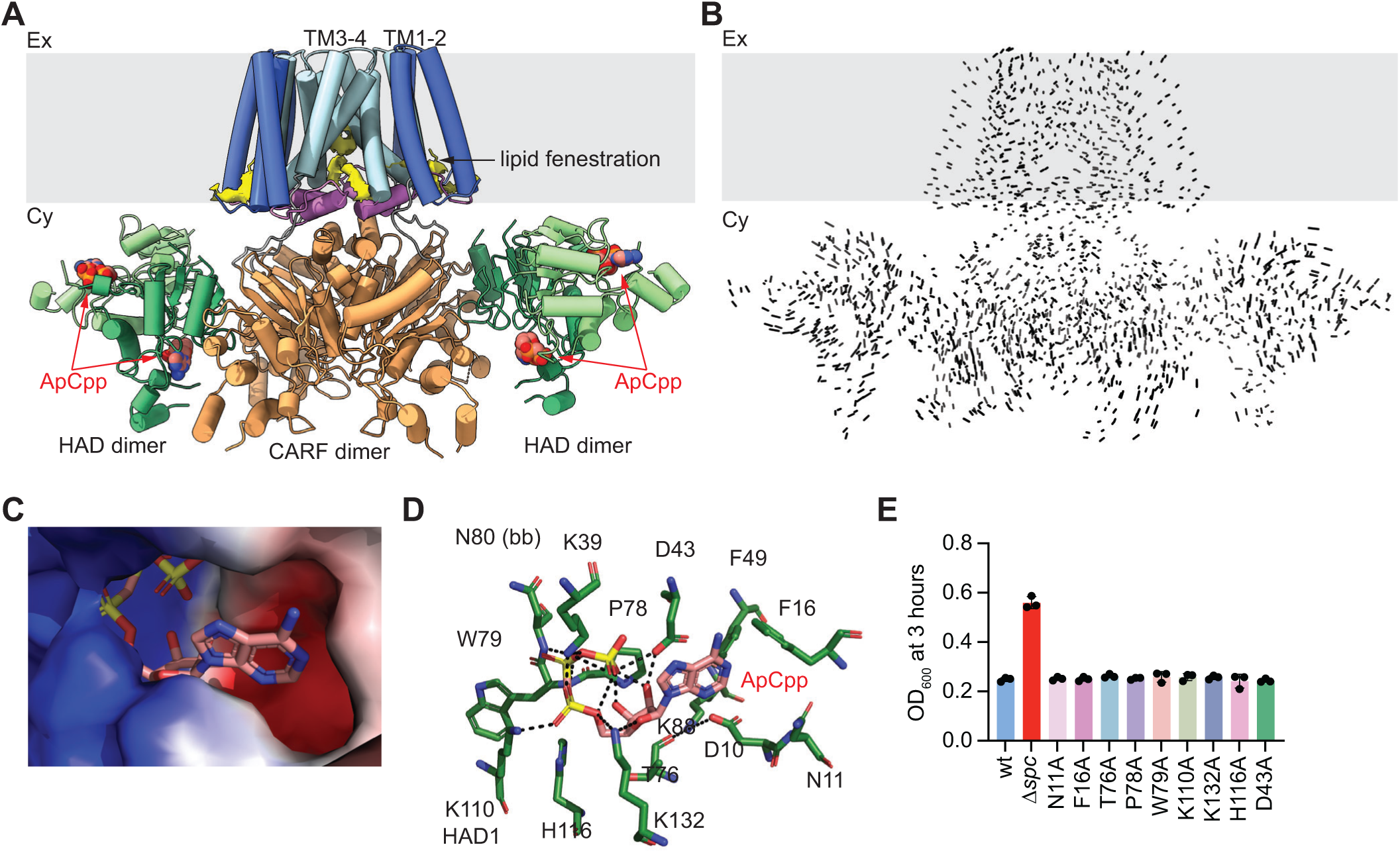
Binding of ApCpp to the HAD domain does not alter Chp1 structure. **(A)** Cryo-EM map of Chp1 in the presence of the ATP analog ApCpp, displaying a tetrameric organization with both the HAD and CARF domains located in the cytosol, while the TM domain anchors the protein to the membrane region (grey area). Lipid fenestrations are shown in yellow in the TM domain. The four ApCpp molecules, each bound to a HAD domain, are shown in red. **(B)** Superposition of apo-Chp1 and ApCpp-bound Chp1 structures showing RMSD of 1.4 Å. **(C)** Electrostatic surface representation of the ATP binding pocket with bound ApCpp. **(D)** Residues lining the ATP binding site of the HAD domain. (bb) indicates backbone interaction. **(E)** Growth of staphylococci carrying pTarget and pCRISPR variants harboring alanine substitutions of Chp1 residues shown in **(D)**, measured as the OD_600_ value 3 hours after addition of aTc. Data are mean of three biological replicates ± s.e.m.

Electrostatic analysis of ApCpp binding pocket revealed a positively charged region that surrounds the phosphate groups and a neutral surface that interacts with the ribose and adenine base (**Fig. 2C**). In this pocket, residues D10, N11, F16, K39, D43, F49, T76, P78, W79, N80, K88, H116, and K132 from one HAD monomer, together with K110 from an adjacent monomer, make contacts with the ApCpp substrate (**Fig. 2D**). To determine the importance of this pocket for Chp1 activity in vivo, we generated the alanine substitutions N11A, F16A, D43A, T76A, P78A, W79A, H116A, and K132A and K110A and tested Chp1-mediated toxicity in staphylococci upon activation of the type III-A CRISPR-Cas response. We found that none of these mutations reduced the growth arrest caused by Chp1 (**Figs. S3C, 2E**), results that are in line with previous studies that reported the persistence of HAD domain activity upon the change of conserved amino acid residues (*22*).

### Binding of cA_4_ rearranges the cytosolic domains of Chp1

Our previous characterization of Chp1 CARF domain demonstrated that it binds cyclic tetra-adenylate (cA_4_), but neither tri-nor hexa-adenylates(*22*). Therefore, to investigate the molecular mechanism of ligand activation, we determined the cryo-EM structure of Chp1 in complex with cA_4_. Analysis of a single dataset resolved into two distinct cryo-EM reconstructions showing alternative modes of cA_4_ binding, hereafter referred to as class A (**Fig. S4A**) and class B (**Fig. 3A**). Both structures revealed two ligands bound per tetramer, with one cA_4_ molecule occupying a pocket at the interface of each CARF dimer (**Fig. 3B**). This stoichiometry contrasts with the membrane-associated CARF effectors that have been characterized at the structural level, Cap1(*10*) and Csx23(*34*), in which a single cA₄ molecule binds at the tetrameric interface of the CARF-like domain. Beyond displaying a similar ligand binding stoichiometry, class A and B complexes differed significantly when compared to the apo-Chp1 structure. Whereas the class A structure showed minimal conformational differences relative to the apo-Chp1 tetramer, with a root-mean-square deviation (RMSD) of 0.65 Å (**Fig. S4B**), class B complexes exhibited pronounced, opposing lateral motions of the CARF-HAD domains (**Fig. S4D**), not observed for the class A complex (**Fig. S4C**), that resulted in a poor alignment of class B and apo-Chp1 structures with a large RMSD of 3.5 Å (**Fig. S4E**).

**Figure 3.**
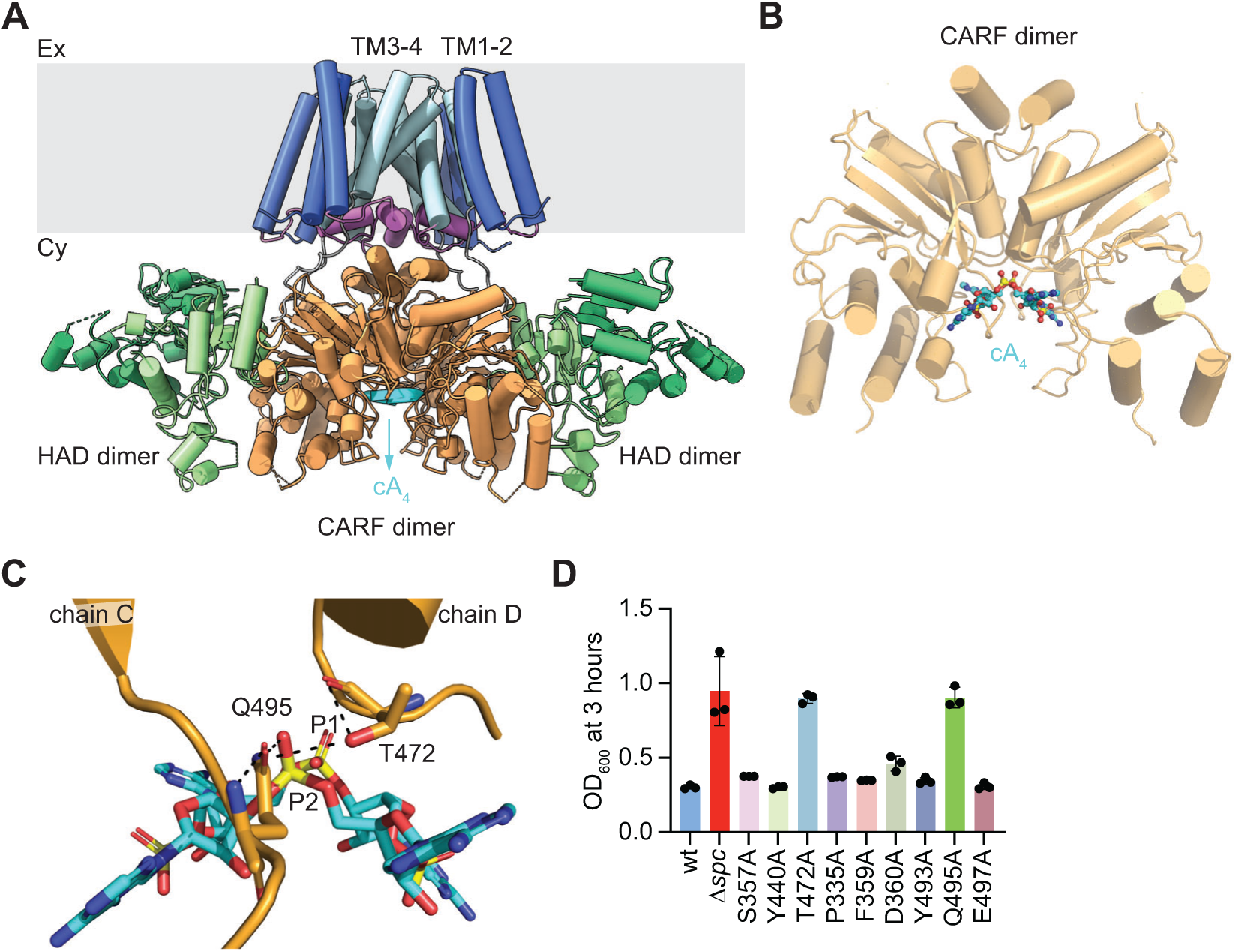
Binding of cA_4_ rearranges the HAD-CARF cytosolic domains. **(A)** Cryo-EM map (class B) of cA_4_-Chp1 displaying a tetrameric organization, with both the HAD and CARF domains located in the cytosol, while the TM domain anchors the protein to the membrane region (grey area). cA_4_ (cyan) binds at the pocket located at the CARF dimer interface**. (B)** CARF dimer highlighting the positioning of concave conformation of the cA_4_ ligand in the class B structure. **(C)** Binding of cA_4_ to the class B CARF dimer ligand pocket showing Q495 (chain C) and T472 (chain D) interaction with the P2 phosphate of cA_4_. **(D)** Growth of staphylococci carrying pTarget and pCRISPR variants harboring alanine substitutions of Chp1 residues shown in **(C)**, measured as the OD_600_ value 3 hours after addition of aTc. Data are mean of three biological replicates ± s.e.m.

The rearrangement of the CARF domain causes very different ligand-binding modes. In the class A tetramer, cA_4_ interacts with the surface of the CARF dimer interface in a convex conformation (**Fig. S4F**). In contrast, in the class B structure binding occurs ∼7.6 Å deeper into the pocket and cA_4_ shifts ∼2 Å to the right, adopting a concave configuration (**Figs. 3B, S4G**). In both conformations, aromatic residues F359, Y440, and Y493 engage in stacking interactions with the adenine bases of cA_4_, and polar and charged residues S318, S357, D360, T472, Q495, and E497 mediate hydrogen-bonding and electrostatic interactions with the base, ribose, and phosphate groups of the ligand (**Figs. 3C, S4H,I**). Two of these residues, Q495 and T472, interact primarily with ribose sugars in the class A structure (**Fig. S4H**), but in the class B CARF domain Q495 shifts ∼3.5 Å to the right and T472 moves ∼4.2 Å upward (**Fig. S4I**), enabling formation of polar contacts with P1 (Q495) and P2 (Q495 and T472) phosphates that stabilize the concave conformation of cA₄ (**Fig. 3C**). We tested the importance of these interactions for Chp1 function in vivo by generating alanine substitutions of residues F359, Y440, Y493, S357, D360, T472, Q495, and E497 and evaluating Chp1 toxicity upon activation of the type III-A CRISPR-Cas response in staphylococci. We found that only the Q495A and T472A mutations, which are engaged in phosphate interactions to stabilize the concave conformation of cA_4_ in the class B CARF domain, abrogated Chp1 function (**Figs. 3D, S4J**). Based on these results, we conclude that the conformational transition of residues Q495 and T472, and the stabilization of the cA_4_ ligand that they mediate, is essential for Chp1 immune function.

### Binding of cA_4_ prevents lipid fenestration

Analysis of the TM domain of the ligand-bound Chp1 complexes revealed a global conservation of this region in both structural classes (**Figs. S5A,B**). Superposition of all four TM helices showed a near-identical alignment, with an RMSD of 0.3 Å, indicating preservation of the overall TM scaffold (**Fig. S5C**). Similarly to the apoChp1 structure, we observed lipid fenestration in the cytosolic half of the TM domain of the class A tetramer (**Figs. 4A,B**). The complex showed the presence of lateral tunnels that connect the surrounding membrane to the central pore (**Fig. 4A**) and are filled with lipid acyl chains that occlude the pore (**Fig. 4B**). In contrast, lipid fenestration was absent from class B complexes (**Fig. 4C**), which showed a pronounced conformational change of the helix within the loop–helix–loop motif of segment J (**Fig. S5B**, black box). This helix rotates 50° toward the central pore (**Fig. 4D**), moves 10 Å deeper into the membrane (**Fig. 4E**) and inserts into the fenestration site to seal the lateral tunnels (**Fig. 4C**). We used Mole2.5 to determine the cavities formed within each Chp1 structural class. This analysis showed a lateral tunnel in the class A structure that forms a continuous hydrophobic conduit with a minimum radius of approximately 2.3 Å (**Figs. 4F,H**), sufficient to accommodate lipid acyl chains. The tunnel is lined by hydrophobic and polar residues from segment J (L231, I235, L236, and L239), TM3 (T251, L252, L254, L255, F257, and V258), TM4 (L292 and I295) and TM1 (F187) (**Fig. S4D**) that create a favorable environment for the penetration and stabilization of membrane lipids. The tunnel entrance, on the other hand, is lined with positively charged residues located on the protein surface that interact with lipid phosphate groups (**Fig. S5E**, R232 and R248). Mutation of each of these two residues to alanine, however, did not affect Chp1 toxicity in vivo (**Figs. S5F,G**). In contrast to the fenestrations observed in class A structure, class B showed a severely constricted tunnel, with a minimum radius of approximately 0.9 Å (**Figs. 4G,H**), that prevents lipid entry. This constriction is a direct result of the inward repositioning of segment J, which occludes the lateral pathway (**Figs. 4D,E,G**). Together, these findings identify segment J as a structural gate that regulates lipid access to the central transmembrane pore of Chp1. While the TM scaffold remains invariant in both structural conformations, segment J undergoes a defined rotational and translational movement that changes the lipid accessibility to Chp1 transmembrane domain.

**Figure 4.**
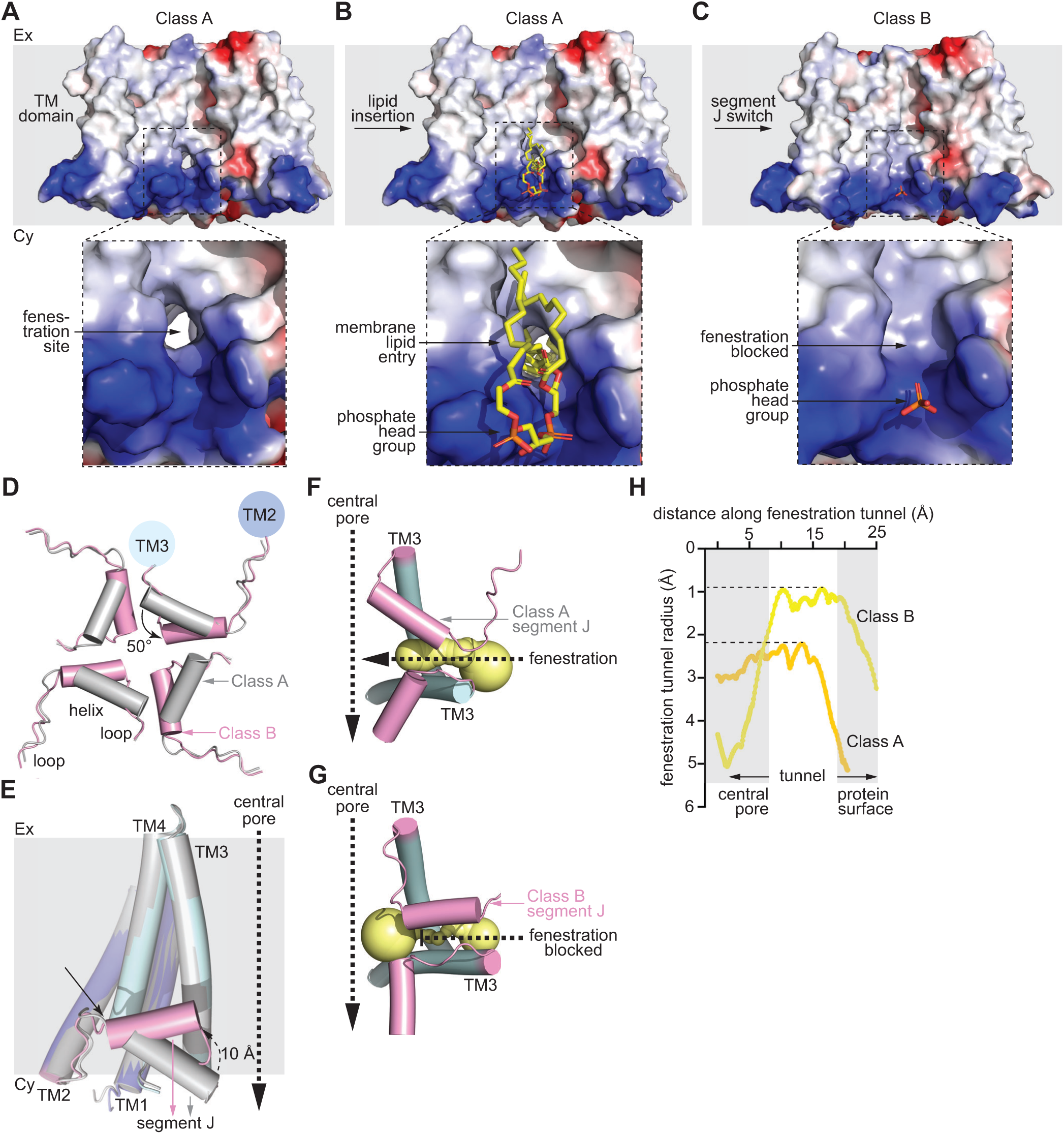
Binding of cA_4_ closes fenestrations within the transmembrane domain. **(A)** Electrostatic surface representation of the TM domain of cA_4_-bound Chp1 class A structure showing a side entry tunnel or fenestration site present in the cytosolic half of the membrane. **(B)** Same as **(A)** but displaying the lipids (yellow) intruding through the fenestration site. The phosphate head group of the lipid interacts with the positively charged (blue) surface of the TM domain while the acyl chain interacts with the side tunnel. **(C)** Same as **(A)** but for the class B structure, showing the block of fenestration site. **(D)** Superposition of segment J from class A (silver) and class B (pink); the latter showing an ∼50° rotation toward the central pore of the helix of the loop–helix–loop motif. **(E)** Side view of the TM domain of subunit 1 of class A (silver) superimposed with that of class B (slate); the latter also showing the ∼10 Å movement of segment J (violet) towards the membrane into the fenestration site. **(F)** Lateral fenestration tunnel analysis by Mole2.5 software for the class A structure. The tunnel is shown in yellow. **(G)** same as **(F)** for class B structure. **(H)** Analysis of the estimated tunnel radius across the fenestration length, measured as the distance in Å from the central pore to the protein surface, for both class A and class B structures, determined using Mole2.5 software.

### Ligand-mediated opening of Chp1 central pore

While lipids that enter through fenestration tunnels occlude the pore formed by the TM helices in both the apoChp1 (**Fig. 1F**) and Chp1-cA_4_ class A (**Fig. 5A**) structures, the Chp1-cA_4_ class B tetramer displayed an open pore that traverses the membrane (**Fig. 5B**). This is because the conformational switch of segment J not only prevents fenestration but also remodels the cytosolic entrance of Chp1 central pore. In the class A structure, segment J lines the cytosolic pore with hydrophobic residues L236, L240, and I244, and therefore generates an environment favorable to lipid occupancy (**Figs. 5C,D**). In contrast, the structural rearrangement of the class B complex reorients residue N242 of segment J toward the cytosolic entrance of the pore, converting the local surface from hydrophobic to polar (**Fig. 5E**).

**Figure 5.**
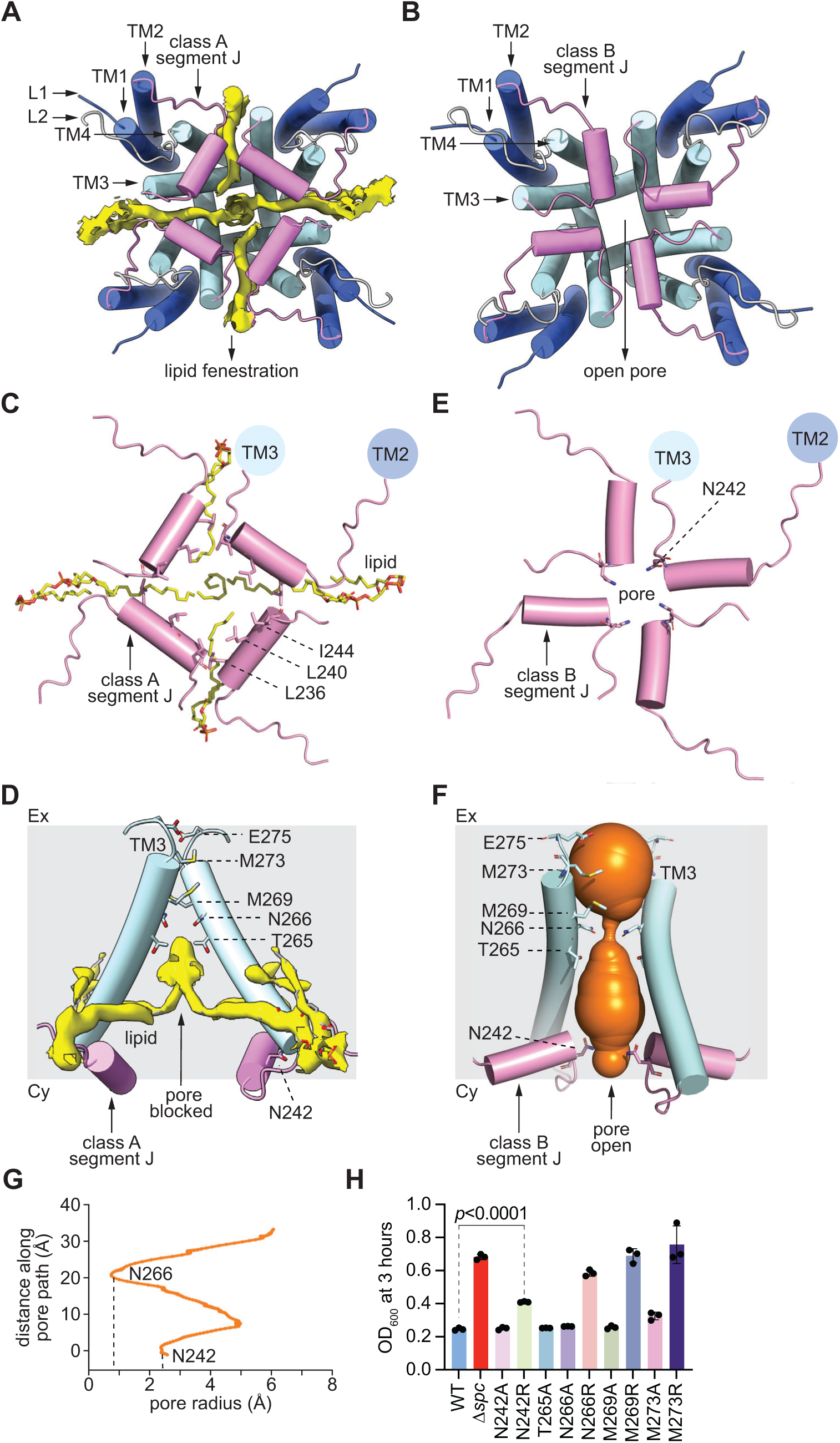
Binding of cA_4_ opens the central pore. **(A)** Cytosolic view of the TM domain showing four lateral lipid fenestrations blocking the central pore in cA_4_-bound Chp1 class A structure. **(B)** Same as **(A)** for the class B structure, highlighting absence of lipid fenestrations, which results in the unblocking of the central pore. **(C)** Cytosolic view of segment J positioning in the class A structure, highlighting the hydrophobic residues lining the cytosolic entrance of the central pore. **(D)** Side view of the TM domain in the class A structure showing lipids that occlude the central pore. The pore-lining residues are noted. **(E)** Same as **(C)** for the class B structure, displaying the cytosolic pore entrance lined by the polar residue N242. **(F)** Analysis of the estimated pore radius across the pore length, measured as the distance in Å from the cytosolic to the extracellular sides of the pore, determined using Mole2.5 software. The graph highlights a constricted section lined by residue N266 and the cytosolic entrance of the pore lined by residue N242. **(H)** Growth of staphylococci carrying pTarget and pCRISPR variants harboring alanine or arginine substitutions of Chp1 residues shown in **(F)**, measured as the OD_600_ value 3 hours after addition of aTc. Data are mean of three biological replicates ± s.e.m.

We analyzed the open pore formed by the TM3 helices and segment J using Mole2.5. We found a constriction of the transmembrane channel formed by four circularly arranged N266 residues (**Fig. 5F**) that reduces the radius to ∼0.8 Å, most likely generating an ion selectivity filter (**Fig. 5G**). Other pore-lining polar residues include N242 from segment J mentioned above, E275 located at the extracellular opening of the pore, and T265 from TM3, all of which could support the formation of a putative cation-conducting pathway (**Fig. 5F**). In addition, M269 and M273, also from TM3, cover the inside of the pore. We tested the importance of these residues for Chp1 function in vivo through mutation to alanine, a neutral substitution, or to arginine, to assess the impact of a bulkier, positively charged side chain that should interfere with the transport of a cation substrate. In all cases, arginine, but not alanine, substitutions, limited the cell toxicity of activated Chp1 (**Figs. 5H, S5H**), a result that suggests that pore occlusion affects the function of this CARF effector. Interestingly, it was not possible to obtain a T265R mutant, potentially due to the generation of a dysregulated pore in the absence of cA_4_. Altogether, these results support a model in which the cA_4_-induced conformational switch of segment J simultaneously seals the lateral fenestrations, converts the cytosolic pore entrance from hydrophobic to negatively polar, and prevents the occlusion of the pore by lipid acyl chains, effectively opening the pore and establishing a plausible route for ion permeation through the TM domain of Chp1.

### Ligand binding does not alter ATP recognition by the HAD domain

To investigate the effects of cA₄ binding on the interactions of the HAD domain with ATP, we determined the cryo-EM structure of Chp1 in the presence of both the ligand and the substrate analog ApCpp. We obtained two structural classes (A, **Fig. S6A**; B, **Fig.S6B**) corresponding to the previously described modes of cA₄ binding in the absence of ApCpp, both showing stable binding of the ATP analog. The class A structure aligned closely with apo-Chp1 (RMSD 1.6 Å; **Fig. S6C**), whereas comparison of class A to B complexes revealed substantial cytosolic domain motions (RMSD 3.1 Å; **Fig. S6D**), consistent with our comparison of cA_4_-bound class A and B structures obtained in the absence of the ApCpp substrate (**Fig. S5B**). These structural rearrangements, however, involved the previously observed inward repositioning of segment J (**Figs. 4D,E**) and the opposing lateral motions of the CARF-HAD domains (**Fig. S4D**), but not the ATP binding pocket. Indeed, superposition of ApCpp-bound HAD domains from both structural classes showed a minimal deviation (RMSD 0.74 Å, **Fig. S6E**), demonstrating that cA₄ binding does not alter the interaction between the HAD domain and its substrate. Therefore, these findings define the HAD substrate-binding site as a preformed composite pocket that engages ATP without structural rearrangements, revealing a rigid mode of nucleotide recognition within the tetrameric Chp1 assembly.

### Chp1 causes membrane depolarization during the Type III CRISPR-Cas response

The structural characterization of Chp1 revealed that activation by cA_4_ does not significantly affect the interaction of the HAD domain with its nucleotide substrate, but instead remodels the TM domain to clear fenestrations from intruding lipids and open a transmembrane pore. Structural data was supported by mutational analysis of Chp1, which demonstrated that changes of key residues of the TM domain and segment J, but not of those that form the ATP binding pocket of the HAD domain, perturbed Chp1 toxicity in vivo. These results therefore contradict our previous conclusion that this activity is caused by dephosphorylation of ATP and dATP, which was based on the measurement of nucleotides in cell extracts upon activation of Chp1 in staphylococci, and in vitro experiments with the purified HAD domain(*22*). In these experiments, the challenges of purifying the full-length protein, however, prevented us from testing whether dephosphorylation depended on binding cA_4_, as it should be expected. We therefore incubated purified full-length Chp1 (2 µM) with ATP or dATP (1 mM) and Mn^2+^ (1 mM), in the presence or absence of cA_4_ (10 mM), and analyzed the reaction products using HPLC. We detected dephosphorylation products for both nucleotide substrates, but the extent of this activity did not change significantly in the presence of the cyclic nucleotide signal. Generation of ADP from ATP was subject to a non-significant increase of 36 % to 40 % in the presence of cA_4_ (**Fig. 6A**), and conversion of dATP to dADP occurred at a 40 % rate irrespective of the presence or absence of ligand (**Fig. 6B**). Moreover, assays with purified F16A and D43A Chp1 mutants, residues involved in ApCpp binding that are not important for Chp1-mediated toxicity in vivo (**Figs. 2D,E**), did not change the extent of ATP and dATP dephosphorylation (**Figs. 6A,B**). Together, these structural, mutational and enzymatic analyses of the HAD domain establish that this domain is not activated during the type III-A CRISPR-Cas response in staphylococci.

**Figure 6.**
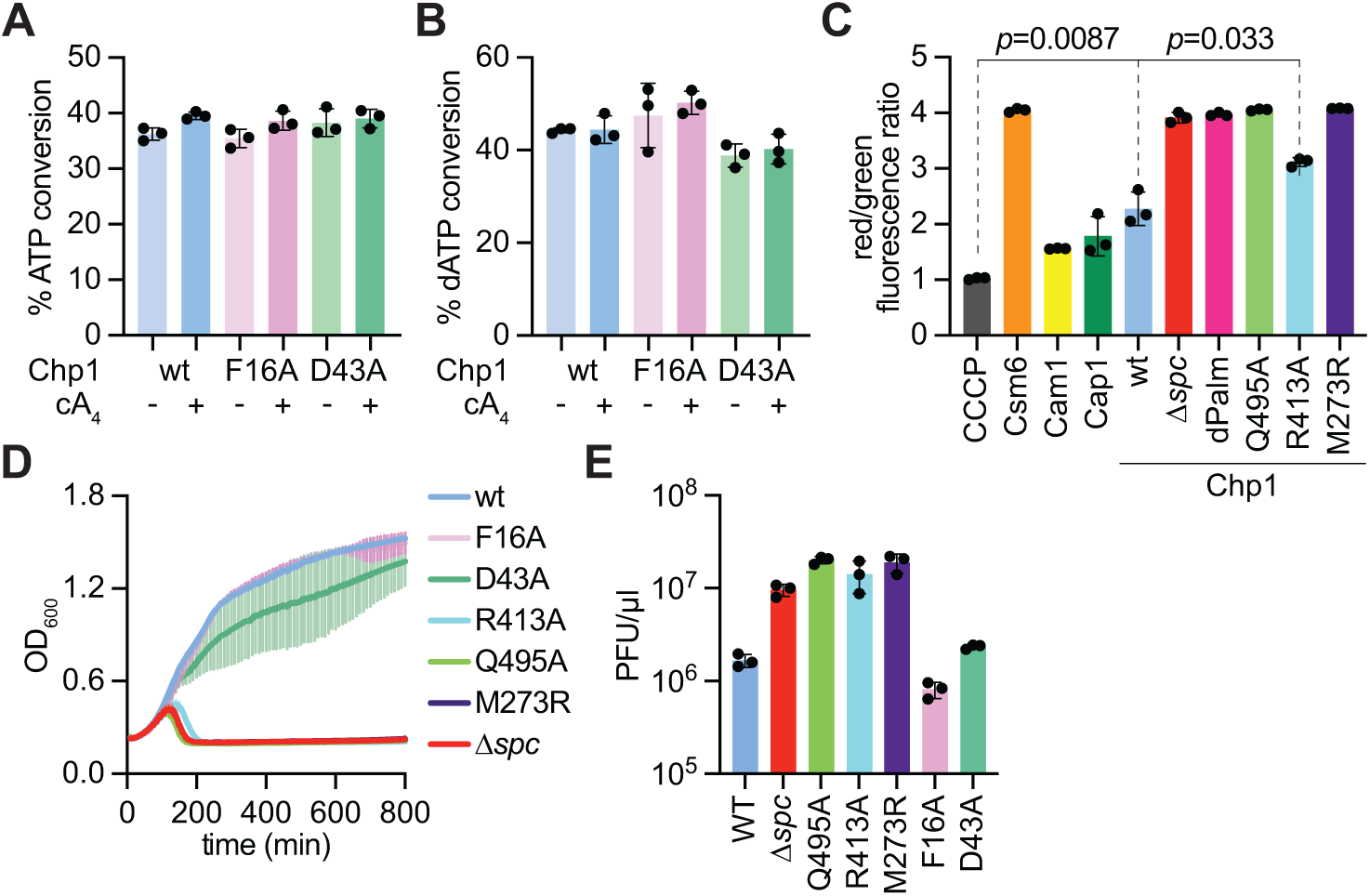
Chp1 activation causes membrane depolarization in staphylococci. **(A)** Quantification of the percent of ATP substrate (1 mM) converted to ADP by purified Chp1 (wild-type and mutant versions; 2 μM) with Mn^+2^ (1 mM) in the presence or absence of cA_4_ (10 mM). Reactions were performed in triplicate, and curve heights for ATP and ADP peaks were used to determine the % conversion. **(B)** Same as **(A)** but using dATP as a substrate. **(C)** Quantification of red/green fluorescence intensity ratio (488B/488C channels) obtained using flow cytometry of staphylococci carrying pTarget and different pCRISPR variants stained with DiOC2(3) after induction of cA_4_ synthesis by addition of aTc to the bacterial cultures. Data are mean of three biological replicates +/− s.e.m. *p* values were obtained with two-sided t-tests with Welch’s correction. **(D)** Growth of staphylococci carrying pCRISPR variants harboring substitutions in key Chp1 residues, measured as OD_600_ after infection with φNM1γ6 phage at an MOI of 1. Data are mean of two or three biological replicates ± SEM. **(E)** Enumeration of PFU within staphylococcal cultures harboring the pCRISPR constructs analyzed in **(D)** with a spacer targeting the *gp43* transcript of φNM1γ6 phage. Cultures were collected at 3 hours after infection at an MOI of 1. The mean of two or three biological replicates, ± SEM, is reported.

The opening of the pore observed in the cA_4_-Chp1 structure uncovered a second possibility, which we considered in our initial characterization of Chp1 given the limitations of the study (*22*): that activation of this CARF effector causes membrane depolarization, similar to the case for Cam1 (*18*), Csx23 (*17*) and Cap1 (*19*). To test this, we triggered the synthesis of cOA ligands in staphylococci harboring pTarget and pCRISPR through the addition of aTc to the cultures. We collected cells 30 minutes after induction and monitored membrane potential using the indicator dye 3,3’-dietyloxacarbocyanine iodide (DiOC_2_(3)). In cells with intact membranes DiOC_2_(3) emits both red and green fluorescence; however, membrane depolarization leads to a decrease in the emission of red fluorescence and therefore a reduction of the red/green fluorescence intensity ratio(*35*). As a positive control, we treated staphylococci with carbonyl cyanide m-chlorophenylhydrazone (CCCP), a membrane potential disruptor. In addition to pCRISPR plasmids expressing Chp1, we also tested Cam1 and Cap1 as controls that cause membrane depolarization (*18*, *19*), and Csm6 (*23*) as a control for RNase toxicity not related to membrane depolarization (**Fig. S2C**). Flow cytometry analysis showed that, as expected from our own previous findings, activation of Cam1 (*18*) and Cap1 (*19*), but not Csm6 (*23*), resulted in low red/green fluorescence intensity ratios, similar to the CCCP control levels (**Fig. 6C**). Chp1-expressing staphylococci treated with aTc also showed a low ratio, similar to that of the Cam1 and Cap1 controls, which increased in the absence of a targeting spacer (Δ*spc*) and in the presence of a mutation in *cas10* that inactivates the Palm domain (*cas10*^palm^), i.e., when cA_4_ is not produced by the Cas10 complex (**Fig. 6C**). Mutation of key residues involved in the conformational change of Chp1 required for opening of the transmembrane pore phenocopied the Δ*spc* and *cas10*^palm^ negative controls and prevented depolarization (**Fig. 6C**). These include R413A, a mutation that destabilizes the CARF-HAD surface (**Figs. 1D,E**); Q495A, which disrupts stabilization of cA_4_ binding in the concave configuration (**Figs. 3C,D**); and the M273R substitution that alters the inner surface of the pore formed by the TM3 helices (**Figs. 5F,H**).

Finally, we tested the effect of mutations of residues from different domains in the defense provided by Chp1 against the staphylococcal phage φNM1γ6 (*36*). We measured culture survival following OD_600_ over time after infection, as well as phage propagation through the enumeration of φNM1γ6 plaque-forming units (PFU) on lawns of staphylococci harboring mutated pCRISPR(Chp1) plasmids. We found that the R413A, Q495A and M273R mutants prevented membrane depolarization (**Fig. 6C**) growth arrest upon activation of the type III-A CRISPR-Cas response (**Figs. 1E, 3D, 5H**), and cell survival (**Fig. 6D**) and PFU reduction (**Fig. 6E**) after infection. In contrast, F16A and D43A alanine substitutions, mutations in the ATP binding pocket of the HAD domain that did not affect Chp1 toxicity (**Figs. 2D,E**), nor anti-phage immunity (**Figs. 6D,E**). These results indicate Chp1 activation during the type III-A CRISPR-Cas response causes membrane depolarization, which is most likely responsible for the growth arrest that prevents viral propagation in staphylococci.

## DISCUSSION

Here we characterized the immune mechanism of Chp1, a CARF effector activated during the type III CRISPR-Cas anti-phage response, and found that it provides immunity by causing the depolarization of the host membrane through the formation and opening of a lipid-gated pore. Cryo-EM data revealed that Chp1 assembles into a tetrameric complex with asymmetric cytosolic CARF-HAD domains and a stable transmembrane region that forms a central pore. Membrane-accessible, hydrophobic lateral tunnels, also known as fenestrations, allow the infiltration of lipid acyl chains from the membrane that occlude this pore in the inactive state of Chp1. Upon RNA-guided recognition of viral transcripts the Cas10 complex synthesizes cyclic tetra-adenylate second messengers that bind the Chp1 CARF domain. Ligand binding induces opposing motions between the two cytosolic CARF-HAD dimers, relaying an allosteric signal to the transmembrane region via rearrangement of segment J, a linker in between TM2 and TM3 helices. This transition occludes lateral tunnels, blocks lipid fenestrations, and opens the central pore to enable ion conduction and membrane depolarization, which in turn generates an inhospitable host unable to support viral replication.

Membrane depolarization is highly toxic to the staphylococcal host, and causes a decrease in the concentration of ATP and IMP in vivo, measured by HPLC-coupled high-resolution mass spectrometry in our initial characterization of Chp1 (*22*). Two lines of evidence, biochemical and structural, indicate that the HAD domain may not substantially contribute to nucleotide depletion upon Chp1 activation in vivo. First, purified full-length Chp1 exhibited limited phosphatase activity against ATP and dATP, which did not change in the presence of cA_4_. Second, the ATP pocket within the HAD domain undergoes no detectable structural rearrangement upon ligand binding. These two results are in line with our previous characterization of Chp1’s HAD domain in isolation (*22*). We found both a comparable dephosphorylation enzymatic activity and a similar domain architecture for the AlphaFold 3 model, with an RMSD of 0.673 Å between both structures (**Fig. S6F**), including the nucleotide binding pocket (**Fig. S6G**). The main difference is the presence of a coordinated Mn²⁺ ion within the active site in the AlphaFold 3 model, for which we did not observe a corresponding metal Coulomb potential in the cryo-EM map. The consistency of these results for the full-length protein in the presence or absence of ligand, and for the isolated HAD module, a condition that excludes any effect of cA_4_ on this domain, indicate that activation of Chp1 in vivo cannot trigger growth arrest through nucleotide dephosphorylation, as we previously concluded. This conclusion is supported by the inability to disrupt Chp1 activity through the introduction of mutations in residues that are located in the HAD phosphatase conserved motifs of Chp1 (*22*), nor in those that interact with the ATP analog ApCpp. On the other hand, the lack of such inactivating mutations prevent us from unequivocally concluding about the role of dephosphorylation in Chp1 activity. Given that only the complete deletion of the HAD domain abolished cellular toxicity and anti-phage immunity (*22*), we propose that this region of Chp1 serves a structural role necessary for the allosteric conformational change induced by ligand binding.

If cA_4_ does not increase the enzymatic activity of the HAD domain, then the reduction in ATP and IMP intracellular concentration previously observed upon Chp1 activation in vivo (*22*) could be a consequence of membrane depolarization. The proton motive force (PMF) generated across the cell membrane through respiration is utilized to generate ATP via ADP phosphorylation by the F_1_F_0_-ATPase in both mitochondria and bacteria (*37*), including staphylococci (*38*). Therefore, it is conceivable that the disruption of the PMF caused by membrane depolarization during Chp1 activation severely affects ATP levels. In the absence of F_1_F_0_-ATPase activity, cells would employ biosynthetic pathways to restore the ATP concentration. Since IMP serves as the central branch-point precursor for purine nucleotide biosynthesis, giving rise to both AMP and GMP (*39*), it is possible that this nucleotide is also depleted upon membrane depolarization Additional experimentation will be required to test this hypothesis and more generally to identify a mechanistic link between the membrane depolarization and nucleotide depletion phenotypes observed upon Chp1 activation.

Previous work described three transmembrane CARF effectors - Cam1, Csx23, and Cap1 - which mediate growth arrest upon activation by viral infection. Cam1 causes membrane depolarization and is predicted to form a pore, but neither its structure nor the mechanism by which it is activated upon cOA binding could be solved(*9*). Similarly, the structure and the molecular mechanism of activation of full-length Csx23 has yet to be determined (*34*). Pulse Dipolar Electron Paramagnetic Resonance Spectroscopy revealed that cA_4_ binding causes a conformational change in Csx23 that is consistent with a shift in the transmembrane helices, a result that suggests that this CARF effector also causes membrane disruption. In contrast to Cam1 and Csx23, Cap1 structure and mode of activation has been characterized in great detail (*10*). It was shown that cA_4_ binding leads to a conformational change of the transmembrane region that causes the widening of an otherwise constricted pore, leading to membrane depolarization and cell toxicity. Unlike Cap1, Chp1 transmembrane helices show minimal conformational variability across the apo or cA_4_-bound forms, indicating that pore activation is not regulated by large-scale remodeling of this region. Instead, Chp1 is autoinhibited in the resting state through lipids that penetrate the pore through fenestrations, which are closed by ligand activation to block lipid intrusion and open the pore. This mechanism enables dynamic control of pore opening without major conformational changes of the transmembrane core.

Hydrophobic fenestrations have been widely documented in voltage-gated ion channels (*24*, *26*–*28*, *30*), TRP channels(*32*), and two-pore channels (*25*, *33*). In humans, a lipid-gated pore activation resembling the Chp1 mechanism has been observed in calcium-activated potassium channels (BK channels), where the pore remains occluded in the Ca^2+^ −free state by lipids entering via fenestrations, and Ca^2+^ binding induces fenestration closure, lipid release, and ion passage (*31*). In bacteria, lipid fenestrations have been shown to facilitate the passage of hydrophobic molecules into the cell. For example, the sodium channel NavAb contains lateral fenestrations connecting the lipid phase of the membrane to the central cavity of the pore allow the entry of hydrophobic drugs (*24*, *26*, *30*). The discovery of the mechanism of pore opening of Chp1 reveals (i) the existence of ligand-mediated gating via lipid fenestrations in bacteria, and (ii) a previously unrecognized mode of CARF-membrane effector immune response. Our findings raise the possibility that similar mechanisms may activate other membrane-associated effectors during bacterial defense.

## METHODS

### Bacterial growth

*Staphylococcus aureus* strain RN4220 (*40*) was grown at 37°C in brain heart infusion (BHI) medium supplemented with 10 μg mL−1 chloramphenicol to maintain pCRISPR and 10 μg mL−1 erythromycin to maintain pTarget. 5 μM CaCl2 was supplemented in phage experiments unless indicated otherwise.

### Plasmid cloning

The plasmids and oligonucleotides used in this study are detailed in Supplementary File S1. The amino acid sequence of Chp1 was sourced from NCBI GenBank (contig CADBDX010000006 from Bacteroidales bacterium) and synthesized by Azenta.

### Growth curves

For in vivo Chp1 toxicity assays, biological replicates of RN4220 overnight cultures containing pTarget and pCRISPR are diluted 1:100, outgrown for about 80 min to OD ∼0.3. Cells are then seeded in a 96-well plate. To induce targeting, 125 ng ml−1 of aTc is added to the appropriate wells. Absorbance at 600 nm is then measured every 10 min by a microplate reader.

For in vivo antiphage immunity, cells containing various pCRISPR constructs were launched in triplicate overnight, diluted 1:100, outgrown for about 1 h and normalized for optical density. Phage ΦNM1γ642 was added at MOI 1. Cells were seeded into a 96-well plate, and absorbance at 600 nm is then measured every 10 min by a microplate reader.

### Quantification of phage plaques

To obtain plaque-forming unit (PFU) counts from cultures infected with phage, *S. aureus* cultures containing various pCRISPR constructs were launched overnight, diluted 1:100 and outgrown for about 1 h. Cells were then infected with phage ΦNM1γ642 at an MOI of 1, and an aliquot was taken after 3 hours of incubation. Aliquots are filtered with a 0.45 μm filter and spotted on cells lacking a pCRISPR construct to measure PFU, with 10-fold serial dilutions for every spot in a lane.

### Flow cytometry

For the membrane depolarization studies, biological replicates of RN4220 overnight cultures containing pTarget and pCRISPR are diluted 1:100, outgrown for about 80 min to OD ∼0.3 and normalized to 10^7^ cells ml^−1^ in 1 mL of PBS. These cultures were then treated with either 125 ng ml^−1^ aTc or 1.7 μM CCCP (Thermo Fisher), followed by a 1 hour 37 °C incubation with shaking. Then, 15 μM DiOC2 (3) (Thermo Fisher) was added and samples were incubated in the dark at room temperature for 5 min. Cells were then analyzed on a BD LSR-Fortessa (BD Biosciences) using FACSDiva Software version 8.0.1 with 100,000 post-gating events recorded for each sample. Red/green ratios were calculated using the Ratio Tab of the FACSDiva by dividing the signal of 488B (red) by the signal of 488C (green). Ratios are calculated from uncompensated linear data and are always reported as linear data.

### Heterologous expression and purification of N-term-His_6_-Chp1

Full-length N-terminally His_6_-tagged Chp1 protein was expressed using *E. coli* Rosetta^TM^ 2 (DE3) (Novagen) cells. The secondary culture was grown at 37 °C in terrific broth (TB) media and induced with 1 mM isopropyl β-D-1-thiogalactopyranoside (IPTG) as the OD_600nm_ reached to 1.2-1.5. The induced culture was further grown at 16 °C overnight and the cells were harvested and resuspended using the lysis buffer (500 mM NaCl, 25 mM Hepes pH 8, 2 mM β-Mercaptoethanol and 5 % glycerol). The resuspended culture was supplemented with cOmplete mini, EDTA-free protease inhibitor tablets (Sigma) and the cells were lysed using a sonicator at 50 % amplitude and 1 s on and 2 s off pulse for a total of 20 minutes. The cell debris were separated by a brief centrifugation and the supernatant was used for harvesting the membrane. The proteins were extracted from the membrane using 20 mM DDM (n-Dodecyl β-D Maltopyranoside) (Anatrace) detergent. After separating the undissolved membrane by centrifugation, the supernatant was loaded to a pre-equilibrated 5 ml HisTrap column (Cytiva). The column was extensively washed with the lysis buffer supplemented with 40 mM imidazole and 1 mM DDM. The protein was eluted using the lysis buffer supplemented with 300 mM imidazole and 1 mM DDM. The eluted protein fractions were checked using NuPAGE^TM^ 4-12 % Bis-Tris gel (invitrogen). The pure protein fractions were pulled and concentrated to perform size column chromatography (SEC) using Superose 6 increase 10/300 GL column. The SEC running buffer was 25 mM Hepes pH 8, 200 mM NaCl, 2 mM β-Mercaptoethanol, 5 % glycerol supplemented with 375 µM GDN (glycol-diosgenin, Anatrace) detergent.

### SEC-MALS analysis of N-terminal-His_6_-Chp1

The SEC-MALS analysis was performed with purified N-terminal-His_6_-Chp1 protein present in GDN detergent using Superdex 200 increase 10/300 GL column. The SEC-MALS instrument (Wyatt) which has multi-angle light scattering detector and refractive index detector was connected to an AKTA-Pure UV detector. The SEC buffer was used for the SEC-MALS runs. ASTRA 6 software was used for data analysis using protein conjugate method. The UV signal of the AKTA-Pure instrument was converted to analogue signal with a conversion factor of 1,000 mAU = 1 V. For protein conjugate analysis the refractive index increment (dn/dc) value for the protein was 0.185 ml/g and for the detergent was 0.14 ml/g.

### In vitro dephosphorylation reactions

Dephosphorylation reactions were performed in vitro by incubating substrates at 37°C for 2 hours. Reaction substrates were incubated at a final concentration of 1 mM with 2 µM Chp1 in reaction buffer [25 mM Hepes pH 8, 200 mM NaCl, 2 mM β-Mercaptoethanol, 5 % glycerol, 1mM MnCl_2_ supplemented with 375 µM GDN (glycol-diosgenin, Anatrace) detergent], with or without 10mM cA_4_. After incubation at 37°C, reactions were quenched by heating mixtures to 65°C for 5 min. To remove the GDN, samples were mixed with a 1:1 volume of equilibrated Bio-Beads SM-2 Adsorbent Media (Bio-Rad 1528920) and incubated on an end-over-end rotator at 4°C for 1 hour. The reaction mixtures were removed to the beads and mixed with 100 μL nuclease-free water before being filtered with Amicon® Ultra Centrifugal Filter, 10 kDa MWCO filters to remove proteins before analysis. 10 μL of filtered reaction products were then injected onto an Agilent Bonus-RP, 4.6 × 150 mm, 3.5 μm Rapid Res. C18 column held at 40°C at a flow rate of 1.2 mL per min. The following mobile phase buffer was used in the elution program: 60 mM K2HPO4, 40 mM KH2PO4, pH 7.0. The elution program was as follows: 0 min 100% buffer, 0% ACN; 2 min 95% buffer, 5% ACN; 4 min 80% buffer, 20% ACN; 5.3 min 75% buffer, 25% ACN; and 6 min 100% buffer, 0% ACN. Chromatograms were collected by monitoring absorbance at 254 nm. To determine substrate conversion, integrated peak heights from ATP or dATP peaks were compared to integrated peak heights from ADP or dADP peaks.

### Cryo-EM sample preparation and data collection on apo- and ligand-bound Chp1 complexes

Purified full-length apo-Chp1 was concentrated to 160 µM for grid preparation. The cA_4_–Chp1 complex was prepared by supplementing apo Chp1 (160 µM) with 1 mM cA_4_. The ApCpp–Chp1 complex was prepared by adding 1 mM ApCpp (Jena Bioscience) and 1 mM MnCl₂ to apo Chp1 (160 µM). The ternary cA_4_–ApCpp–Chp1 complex was prepared by supplementing apo Chp1 (160 µM) with 1 mM cA_4_, 1 mM ApCpp, and 1 mM MnCl₂. All the samples were prepared in the presence of 0.5 mM FOM (Fluorinated Octyl Maltoside) detergent to correct the orientation preference problem. Quantifoil Au R (1.2/1.3) grids were used for all sample preparations. The grids were glow discharged for 2 minutes at 15 mA and frozen at 4°C, 86-90 % humidity, 13 s wait time, 3.5 – 4.5 s blot time and 0 blot force using Vitrobot Mark IV (FEI). The apo-Chp1, cA_4_–Chp1 complex and cA_4_–ApCpp–Chp1 complex datasets were collected at the Simons Electron Microscopy Center, as well as at MSKCC using a Krios G4 microscope equipped with a Falcon 4i detector and an energy filter of 10 eV slit width. The pixel size was 0.725 Å, and data were collected using a defocus range of −0.8 to −2.3 µm. Images were acquired in EER mode with a total electron dose of 60 e⁻ Å⁻². The EER unsampling factor was set to 1, and each movie was fractionated into 45 frames. The ApCpp–Chp1 complex dataset was collected at the Simons Electron Microscopy Center, as well as at MSKCC using a Titan Krios G2 microscope equipped with a Gatan K3 direct electron detector. Images were acquired in super-resolution mode with a pixel size of 0.532 Å, using a total electron dose of 52.9 e⁻ Å⁻² distributed over 60 frames per movie.

### Cryo-EM data processing and refinement on apo- and ligand-bound Chp1 complexes

CryoSPARC v4.4.1 was used for data processing (*41*), while Chimera (*42*) and coot (*43*) were used for model building, Phenix real-space refinement program (*44*) was used for the refinement (**Tables S1 and S2**). Representative cryo-EM map densities for different parts of the protein and the ligands are shown in **Fig. S7**.

#### Processing of apo Chp1 dataset

12,582 movies were collected, and Patch Motion Correction job was used to generate the motion corrected micrographs which were then used as input to the Patch CFT Estimation job. Followed by CTF estimation, 4,737,447 particles were picked up using Blob Picker job and 3,894,840 particles were extracted by the Extract From Micrographs job with an extraction box size of 400 pixel. By iterative rounds of 2D classification job, 999,778 particles were chosen (**Figs. S8A, B**). Further, from these set of particles 118,194 particles were used to generate the 3D ab initio model. The best 3D class was reconstructed using 64,422 particles. Next, Hetero Refinement job was performed to select more good particles from 999,778 particles, which selected 244,444 particles (**Fig. S8A**). Next 3D classification job was performed, and two classes, labeled A and B, could be resolved to high resolution containing 119,356 particles and 85,128 particles, respectively (**Fig. S8C**). Non-uniform Refinement job was used to refine the classes at 2.86 Å and 3.16 Å with C1 symmetry (**Fig. S8D**). The resolution was further improved by refining the maps with C2 symmetry to 2.81 Å and 2.96 Å following the standard FSC cutoff value of 0.143 (**Figs. S8D-F**). Local resolution variations across the cryo-EM map were assessed using the Local Resolution Estimation job (**Figs. S8G, H**). Domain models predicted by AlphaFold3(*45*) were employed for initial rigid-body fitting in UCSF Chimera (*42*) followed by manual model adjustment in Coot (*43*). The model was subsequently refined against the cryo-EM density using Phenix real-space refinement (*44*) to optimize geometry and reduce outliers, yielding a model–map correlation coefficient (CC_mask) of 0.86 for class A and 0.84 for class B (**Table S1**).

#### Processing of ApCpp-Chp1 dataset

4,010 movies were collected for ApCpp-Chp1 dataset, and 3,960 movies were motion corrected using Patch Motion Correction job. These 3960 micrographs were used for CTF estimation by Patch CTF Estimation job. Next, Micrograph Denoiser job was used and the denoised micrographs were used to pick particles using the Blob Picker job. 4,305,154 particles were picked by Blob Picker and out of which 3,821,452 particles were extracted by Extract from Micrographs job using 256 pixel box size (**Fig. S9A)**. After iterative 2D Classification job (**Fig. S9B**), 171,163 particles were selected and used as an input to Ab-Initio Reconstruction job. Out of five ab-initio classes one class with 52,975 particles was chosen for the next step. Hetero-Refinement job was used to select more particles for that class and 220,966 particles were selected (**Fig. S9A**). Next 3D classification job was used with these 220,966 particles and one good 3D class having 86,787 particles could be resolved to 3.83 Å (**Figs. S9C, D**). Further C2 symmetry was applied to that class and the resolution was improved to 3.66 Å following the standard FSC cutoff value of 0.143 (**Figs. S9D-F**). The local resolution for different parts of the map was estimated using Local Resolution Estimation job **(Fig. S9G)**. The AlphaFold3 (*45*) predicted models of different domains were used for the rigid body fitting using Chimera (*42*) and coot (*43*) was used for model building. Phenix real-space refinement program (*44*) was used to remove the outliers and refine the model with a model vs. data correlation value (CC mask) of 0.82 (**Table S1**).

#### Processing of cA_4_-Chp1 dataset

11,562 movies were collected for cA_4_-Chp1, and these movies were motion corrected using Patch Motion Correction job. Patch CTF Estimation job was performed for the CTF estimation of 11,562 micrographs. Blob Picker job was used to pick up 2,961,656 particles which were then extracted using 400-pixel box size (**Fig. S10A**). Iterative rounds of 2D Classification job (**Fig. S10B**) were used to select 145,600 particles. These particles were used for Ab-Initio reconstruction job. Out of three ab-initio reconstructions one reconstruction containing 71,024 particles was selected for selecting more good particles using Hetero-Refinement job. The Hetero-Refinement job selected 219,665 particles which were then used as input in the 3D classification job and classified into five classes (**Fig. S10C**). Out of which two classes, class A (92,958 particles) and class B (60,922 particles) were resolved to high resolution; 3.22 Å for class A map and 3.46 Å for the class B map using Non-uniform Refinement job (**Fig. S10D**). Further, C2 symmetry was applied to both the maps which improved the resolution to 2.98 Å and 3.28 Å for class A and class B maps respectively following the standard FSC cutoff value of 0.143 (**Fig. S10D-F**). Map local resolution was determined using the Local Resolution Estimation job (**Figs. S10G, H**). Structural models predicted by AlphaFold3(*45*) were fitted into the cryo-EM density using Chimera (*42*) and subsequently refined through manual model building in Coot (*43*). Final model optimization and outlier removal were performed using the Phenix real-space refinement (*44*), resulting in a model-to-map correlation (CC mask) of 0.85 for class A and 0.87 for class B (**Table S2**).

#### Processing of ApCpp-cA_4_-Chp1 dataset

In the case of ApCpp-cA_4_-Chp1 sample 9,334 movies were collected and motion correction was performed using Patch Motion Correction job. CTF estimation was performed on 9,334 micrographs and Blob Picker job was used to pick up 3,649,002 particles (**Fig. S11A**). Out of these particles 3,008,770 particles were extracted using 400-pixel box size. Iterative 2D Classification job (**Fig. S11B**) was performed, and 137,497 particles were picked to run the Ab-Initio Reconstruction job. Five ab-initio maps were reconstructed and used as input in the Hetero-Refinement job. Using Hetero-Refinement job, 1,748,069 particles were segregated and cleaned to different classes. Selected 365,748 particles were then used for 3D Classification job (**Fig. S11C)**. Further, Remove Duplicate particle job was performed to remove any duplicate particles. Out of the 3D classes two classes could be resolved to high resolution, 2.53 Å for class A (156,296 particles) and 3.03 Å for class B (82,288 particles) using Non-uniform Refinement job (**Fig. S11D**). Next, C2 symmetry was applied to both class A and class B maps and resolution was improved to 2.33 Å for class A and 2.89 Å for class B maps following the standard FSC cutoff value of 0.143 (**Figs. S11D-F**). Local resolution of the maps was estimated using the Local Resolution Estimation job (**Figs. S11G, H)**. AlphaFold3–predicted models (*45*) were docked into the maps using Chimera(*42*) and manually adjusted in Coot (*43*). The models were further refined using Phenix real-space refinement (*44*) to remove outliers, yielding a model–map correlation (CC mask) of 0.85 for class A and 0.82 for class B (**Table S2**).

## SUPPLEMENTARY FIGURE LEGENDS

**Figure S1.**
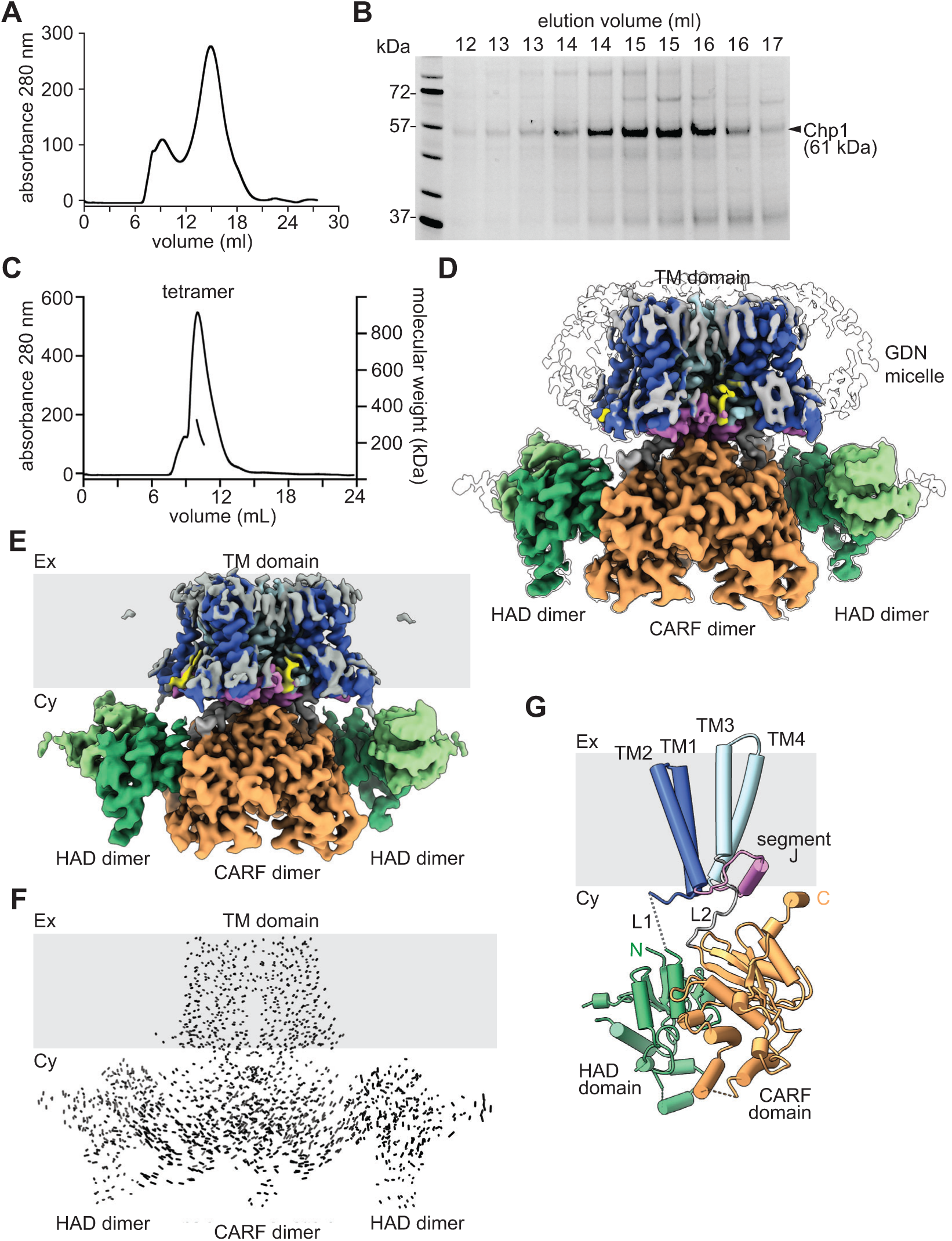
Purification and cryo-EM structure of apo Chp1. **(A)** Size exclusion chromatography (SEC) profile of purified, full-length His_6_-Chp1 protein in GDN detergent. **(B)** SDS-PAGE of the elution fractions collected in **(A)**. **(C)** SEC-MALS analysis with purified N-term His_6_-Chp1 protein reveals tetrameric state (250 kDa ± 2.6 %) in the presence of GDN detergent. **(D)** Cryo-EM map (class A) of apo Chp1 displaying a tetrameric organization, with both the HAD and CARF domains located in the cytosol, while the TM domain anchors the protein to the membrane. The GDN detergent micelle is shown in white, and highlights the boundary of the hydrophobic TM domain and cell membrane. **(E)** Same as **(D)** but for the class B structure. **(F)** Superposition of apo Chp1 class A and B structures; which are highly aligned with an RMSD value of 1.2 Å. **(G)** Structure of Chp1 subunit 1, showing its different structural elements from N- to C-terminal: the HAD domain, flexible linker L1 (dotted line), TM1 entering the membrane (grey area) and turning sharply on the extracellular side to connect with TM2, segment J (a 22-amino acid loop–helix–loop motif), TM3, which crosses the membrane and links to TM4 through a short extracellular loop, linker L2, and the CARF domain.

**Figure S2.**
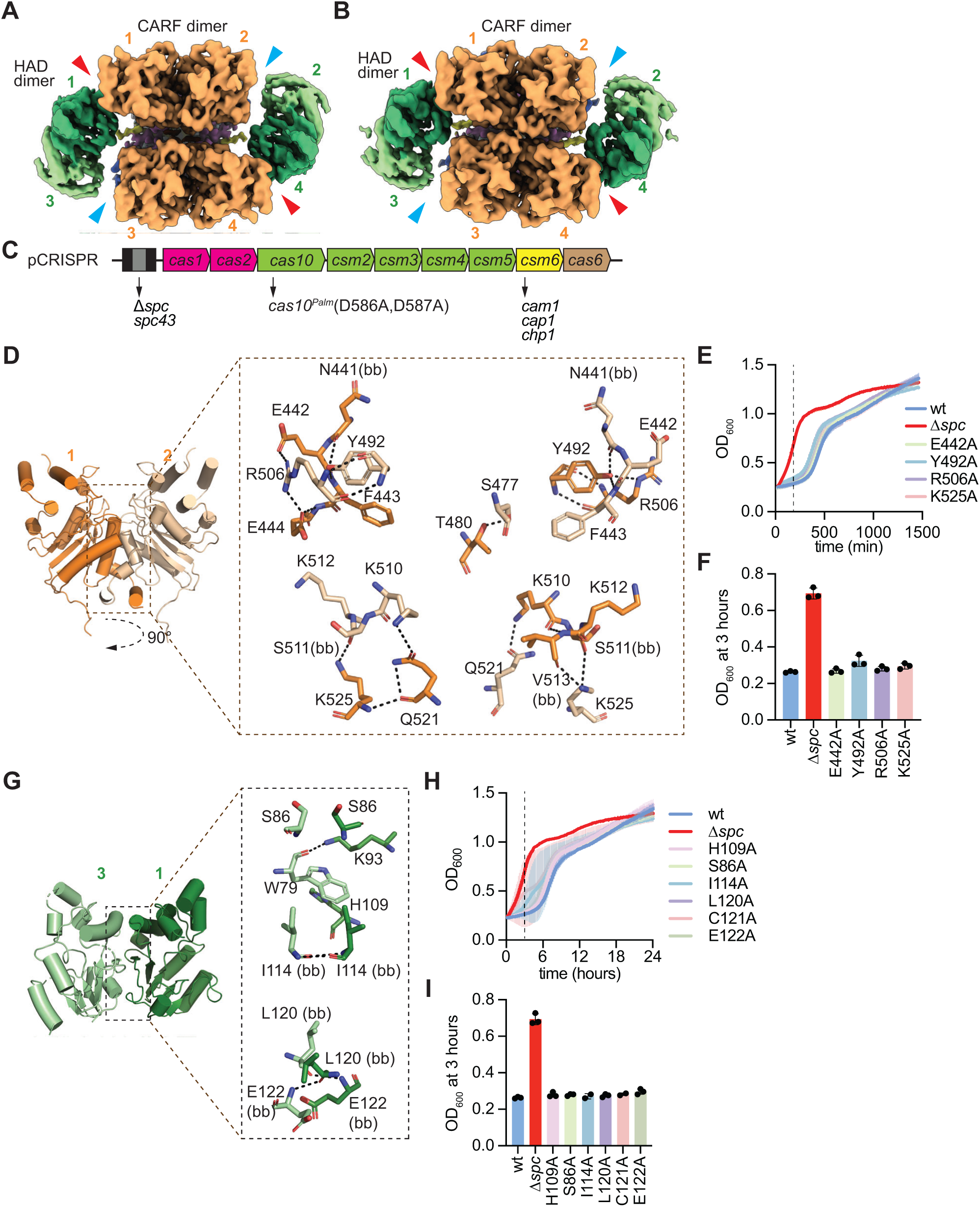
Interactions within CARF and HAD domains in the Chp1 tetramer. **(A)** Cytosolic view of apo Chp1 class A structure showing a CARF-dimer of dimers forming the core of the tetramer and dimeric HAD domains are radially distributed. The HAD and CARF domains interact in subunits 1 and 4 (red arrowheads), whereas in subunits 2 and 3 the HAD domains are positioned farther from the CARF domains (blue arrowheads). **(B)** Same as **(A)** but for the apo Chp1 class B structure. **(C)** Genetic modifications of the *S. epidermidis* RP62a type III-A CRISPR locus cloned into various pCRISPR plasmids. Amino acid substitutions in Cas10 and insertion of different spacer sequences or CARF effectors are indicated. **(D)** CARF dimer interface of subunits 1 and 2 of apo Chp1 class A structure, highlighting the interacting residues. (bb) indicates backbone interaction. **(E)** Growth of staphylococci carrying pTarget and pCRISPR variants harboring alanine substitutions of Chp1 residues shown in **(D)**, measured as the OD_600_ value of the culture after addition of aTc. Data are mean of three biological triplicates ± s.e.m. Dotted line indicates the 3-hour timepoint used for bar graphs. **(F)** OD_600_ value of the cultures shown in **(E)** 3 hours after addition of aTc. Data are mean of three biological replicates ± s.e.m. **(G)** HAD dimer interface of subunits 1 and 3 of apo Chp1 class A structure, highlighting the interacting residues. (bb) indicates backbone interaction. **(H)** Growth of staphylococci carrying pTarget and pCRISPR variants harboring alanine substitutions of Chp1 residues shown in **(G)**, measured as the OD_600_ value of the culture after addition of aTc. Data are mean of three biological replicates ± s.e.m. Dotted line indicates the 3-hour timepoint used for bar graphs. **(I)** OD_600_ value of the cultures shown in **(H)** 3 hours after addition of aTc. Data are mean of three biological replicates ± s.e.m.

**Figure S3.**
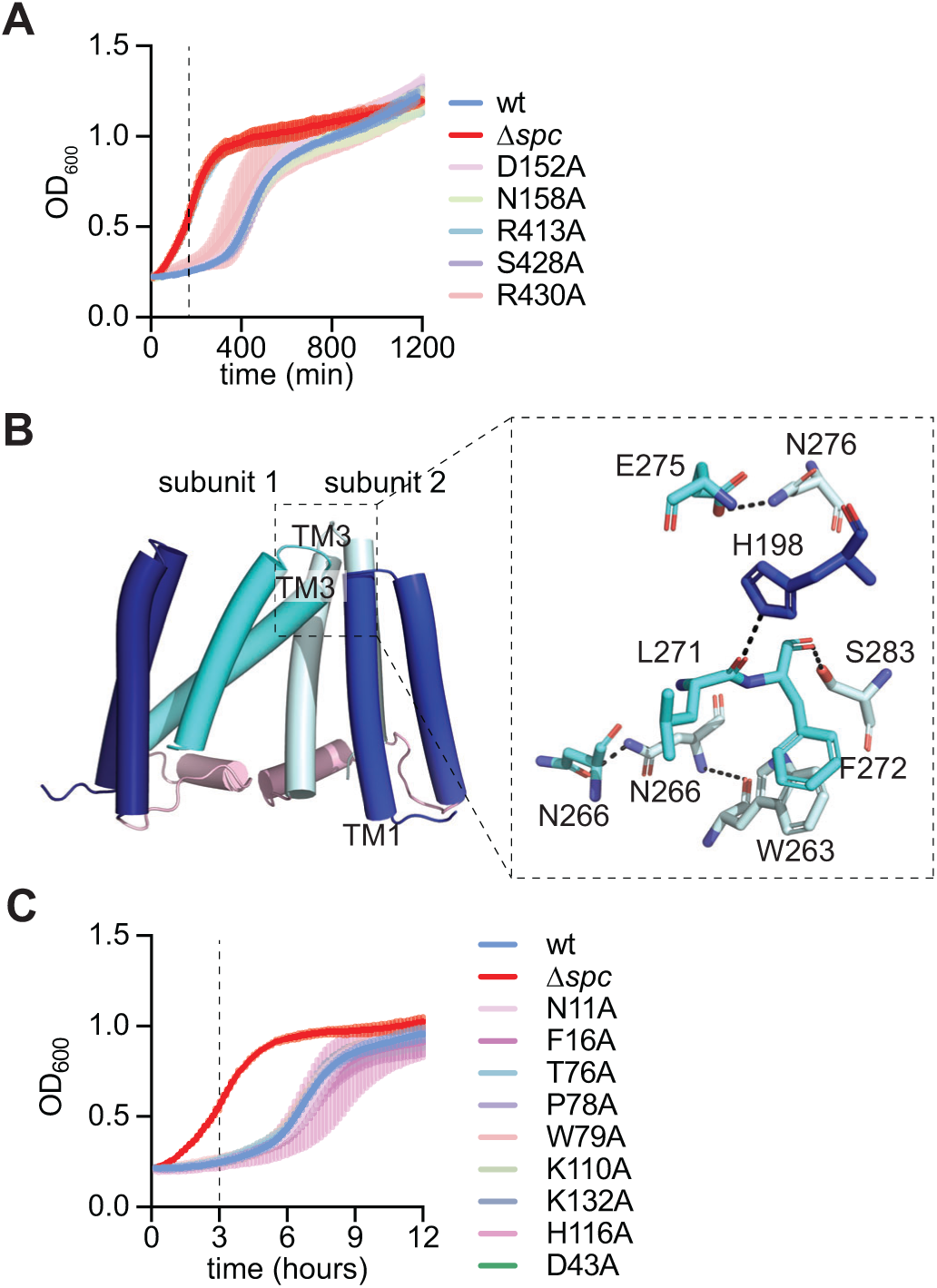
Interactions between the transmembrane helices in the Chp1 tetramer. **(A)** Growth of staphylococci carrying pTarget and pCRISPR variants harboring the indicated alanine substitutions, measured as the OD_600_ value of the culture after addition of aTc. Data are mean of three biological replicates ± s.e.m. Dotted line indicates the 3-hour timepoint used for bar graphs. **(B)** Structure of the tetrameric interface of the TM domain in subunits 1 and 2 of the apo Chp1 class A structure, highlighting the interacting residues. (bb) indicates backbone interaction. **(C)** Growth of staphylococci carrying pTarget and pCRISPR variants harboring the indicated alanine substitutions, measured as the OD_600_ value of the culture after addition of aTc. Data are mean of three biological replicates ± s.e.m. Dotted line indicates the 3-hour timepoint used for bar graphs.

**Figure S4.**
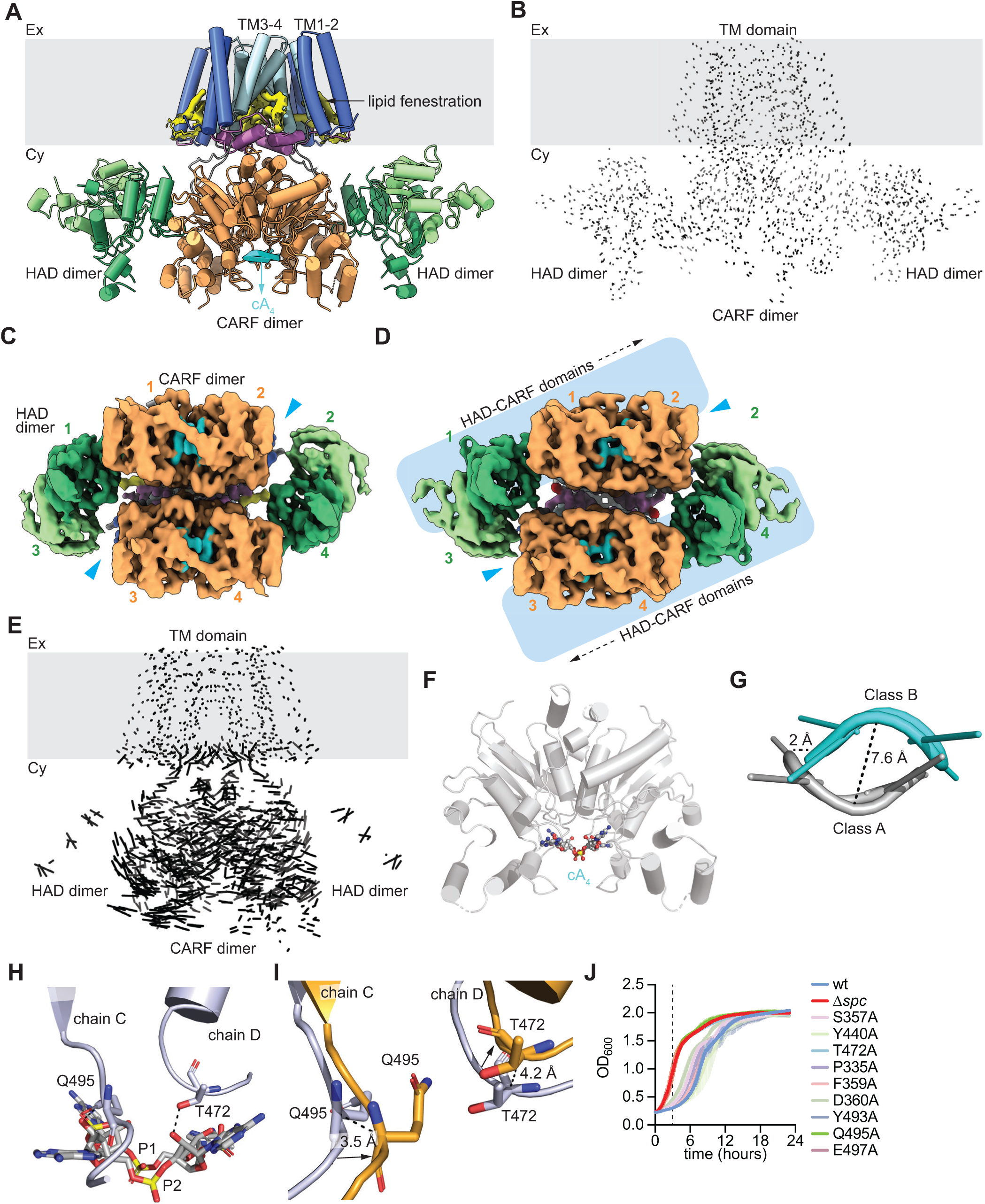
Cryo-EM structure of cA_4_-Chp1. **(A)** Cryo-EM structure (class A) of cA_4_-Chp1 displaying a tetrameric organization, with both the HAD and CARF domains located in the cytosol, while the TM domain anchors the protein to the membrane region (grey area). cA_4_ (cyan) binds at the pocket located at the CARF dimer interface**. (B)** Superposition of apo Chp1 and cA_4_-Chp1 (class A) structures; which are highly aligned with an RMSD value of 0.65 Å. **(C)** Cytosolic view of the cA_4_-bound Chp1 class A cryo-EM map showing two cA_4_ molecules (cyan), with one cA_4_ bound to each CARF dimer. The gaps between the HAD and CARF domains are indicated by blue arrowheads. Orange and green numbers indicate each of the CARF and HAD domains, respectively, of the four Chp1 subunits. **(D)** Same as **(C)** but showing cA_4_-bound Chp1 class B cryo-EM map. The opposite motions of CARF and HAD dimers are indicated by black arrows. **(E)** Superposition of apo Chp1 and cA_4_-Chp1 (class B) structures, which are poorly aligned, especially at the cytosolic domains, with an RMSD value of 3.5 Å. Superposition was limited to the dimerization interface of the HAD domains; the full HAD domains could not be aligned. **(F)** CARF dimer highlighting the positioning of convex conformation of the cA_4_ ligand in the class A structure. **(G)** Superposition of class A (silver, convex) and class B (cyan, concave) cA_4_ binding modes. **(H)** cA_4_-Chp1 class A cA_4_ binding mode, showing Q495 (chain C) and T472 (chain D) interactions with the ligand. **(I)** Superposition of cA_4_ binding pockets for class A (silver) and class B (orange) cA_4_-Chp1 structures, focusing on Q495 (chain C) and T472 (chain D). Black arrows indicate the conformational rearrangement upon transitioning from class A to class B. **(J)** Growth of staphylococci carrying pTarget and pCRISPR variants harboring the indicated alanine substitutions, measured as the OD_600_ value of the culture after addition of aTc. Data are mean of three biological replicates ± s.e.m. Dotted line indicates the 3-hour timepoint used for bar graphs.

**Figure S5.**
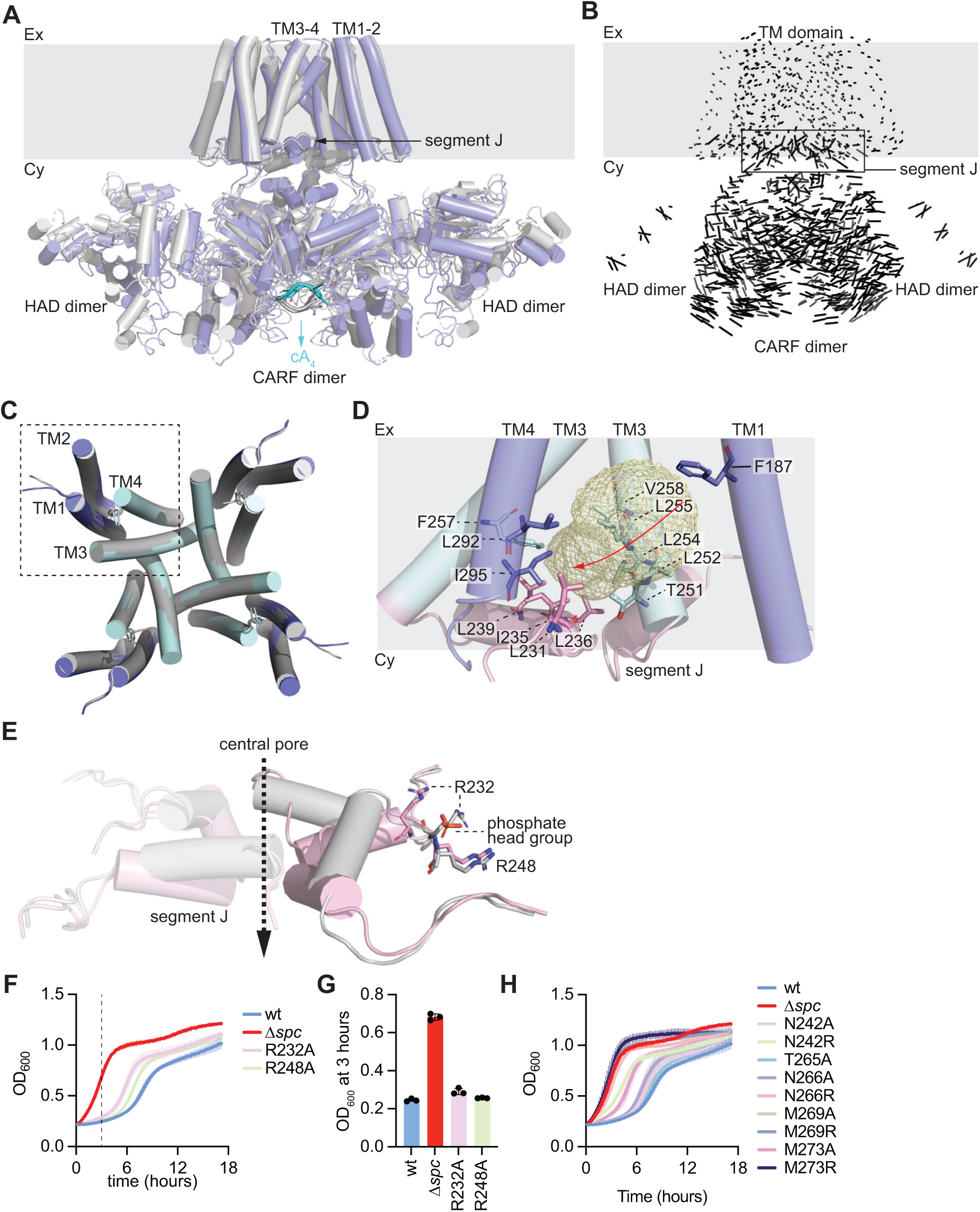
cA_4_ binding releases the lipid block of Chp1 transmembrane pore. **(A)** Superposition of cA_4_-Chp1 class A (silver) and class B (slate) structures, revealing highly alignmed TM domains but large variations in the cytosolic domains and segment J. The change in the conformation of the cA_4_ ligand (cyan, concave in class B; silver, convex in class A) is noted. **(B)** Vector representation of the conformational changes shown in **(A)**. The box highlights the rearrangements of segment J. RMSD 3.4 Å. **(C)** Superposition of class A (silver) and class B (slate) TM domain structures, demonstrating close alignment of all four transmembrane (TM) helices. **(D)** Residues lining the fenestration tunnel (yellow mesh) within the TM domain of cA_4_-Chp1 class A structure. **(E)** Superposition of segment J of cA_4_-Chp1 class A (silver) and class B (violet) structures, showing the R232 and R248 residues at the surface side of the fenestration tunnel, which interact with lipid phosphate head groups. **(F)** Growth of staphylococci carrying pTarget and pCRISPR variants harboring the indicated alanine substitutions, measured as the OD_600_ value of the culture after addition of aTc. Data are mean of three biological replicates ± s.e.m. Dotted line indicates the 3-hour timepoint used for bar graphs. **(G)** OD_600_ value of the cultures shown in **(F)** 3 hours after addition of aTc. Data are mean of three biological replicates ± s.e.m. **(H)** Growth of staphylococci carrying pTarget and pCRISPR variants harboring the indicated alanine or arginine substitutions, measured as the OD_600_ value of the culture after addition of aTc. Data are mean of three biological replicates ± s.e.m. Dotted line indicates the 3-hour timepoint used for bar graphs.

**Figure S6.**
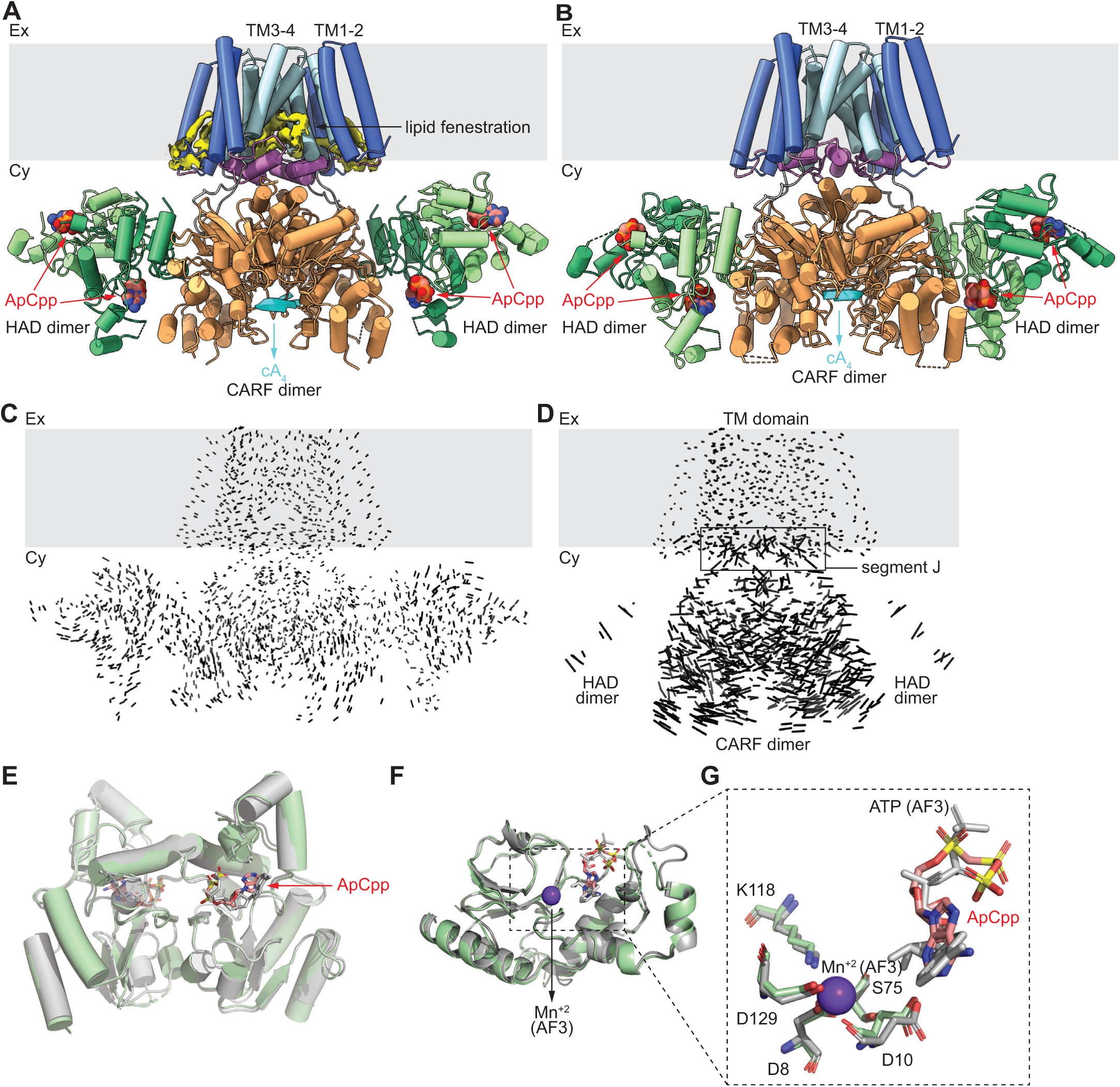
cA_4_ binding does not affect the interactions between the HAD domain and its ATP substrate. **(A)** Cryo-EM map (class A) of cA_4_-Chp1 in the presence of the ATP analog ApCpp (red) displaying a tetrameric organization, with both the HAD and CARF domains located in the cytosol, while the TM domain anchors the protein to the membrane region (grey area). cA_4_ (cyan) binds at the pocket located at the CARF dimer interface**. (B)** Same as **(A)** but for the cA_4_-Chp1-ApCpp class B structure. **(C)** Superposition of apo Chp1 and cA_4_-Chp1-ApCpp (class A) structures, which are highly aligned with an RMSD value of 1.6 Å. **(D)** Same as **(C)** but comparing class A to class B cA_4_-Chp1-ApCpp structures, which are poorly aligned with an RMSD value of 3.1 Å. **(E)** Superposition of the dimeric HAD domains from cA_4_-Chp1-ApCpp class A (silver) and class B (pale green) structures, which are highly aligned with an RMSD value of 0.74 Å. **(F)** Superposition of the HAD domain from an AlphaFold3 model (silver) and cA_4_-Chp1-ApCpp class B structure (pale green), which are well aligned with an RMSD value of 0.67 Å. (**G**) Expanded view of the substrate binding pocket of HAD domain from panel F. The Mn+2 ion was modeled in AlphaFold but the corresponding metal Coulomb potential was not observed in the cryo-EM map.

**Figure S7.**
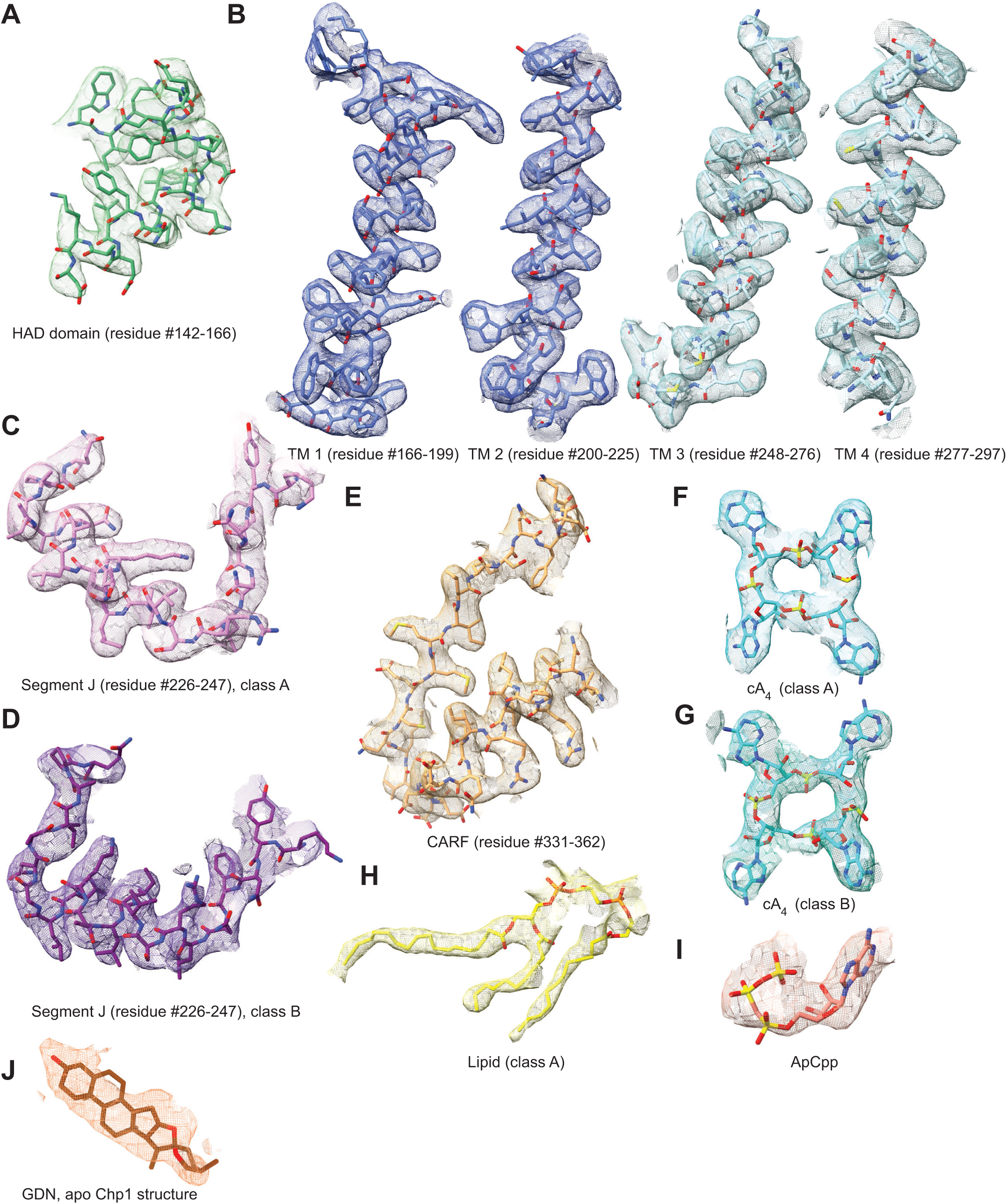
Representative cryo-EM maps for different regions of Chp1 protein and the ligands. **(A)** The map corresponding to the HAD domain residue-number 142-166 is displayed from cA_4_-bound Chp1 class1 structure. **(B)** Similar representation as in panel A, for four TM helices. **(C)** Similar representation as in panel A, displaying segment J for cA_4_-bound Chp1 class A structure. **(D)** Same as in panel C but for cA_4_-bound Chp1 class B structure. **(E)** The map corresponding to the CARF domain residue-number 331-362 is displayed from cA_4_-bound Chp1 class A structure. **(F)** The map corresponding to cA_4_ molecule from cA_4_-bound Chp1 class A structure is displayed. **(G)** Same as panel F, but from cA_4_-bound Chp1 class B structure. **(H)** The map corresponding to the lipid molecule from cA_4_-bound Chp1 class A structure is shown. **(I)** The map corresponding to ApCpp ligand is shown from the ApCpp-cA_4_-bound Chp1 class A structure. **(J)** The map corresponds to GDN detergent from apo Chp1 class A structure. All the maps are displayed at around contour level of 3.9 RMSD (coot) value.

**Figure S8.**
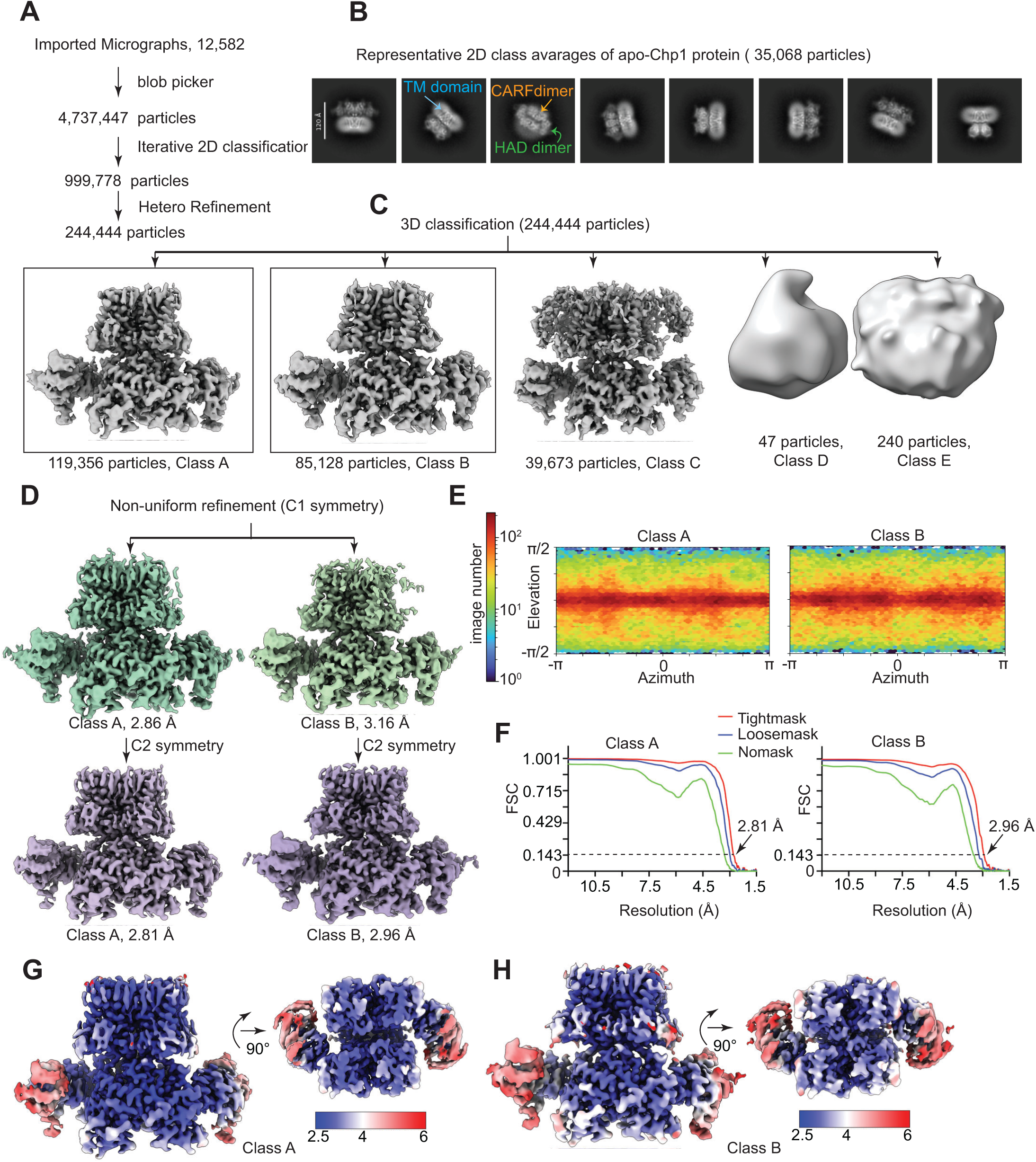
Cryo-EM data processing workflow for apo Chp1. **(A)** Overview of the particle selection workflow. **(B)** Representative 2D class averages of apo Chp1, revealing clear structural features corresponding to the CARF, HAD, and transmembrane (TM) domains. **(C)** 3D classification of the selected particles yielded five distinct classes. Class A and Class B (highlighted by black boxes) were selected for subsequent analysis. **(D)** The reconstructed 3D maps were further refined using non-uniform refinement. **(E)** Angular distribution of particles contributing to the final reconstruction. **(F)** Fourier shell correlation (FSC) curves from the non-uniform refinement job, calculated using no mask, a loose mask, and a tight mask. **(G-H)** Local resolution estimates for Class A and Class B. Resolution values are shown in angstroms (Å).

**Figure S9.**
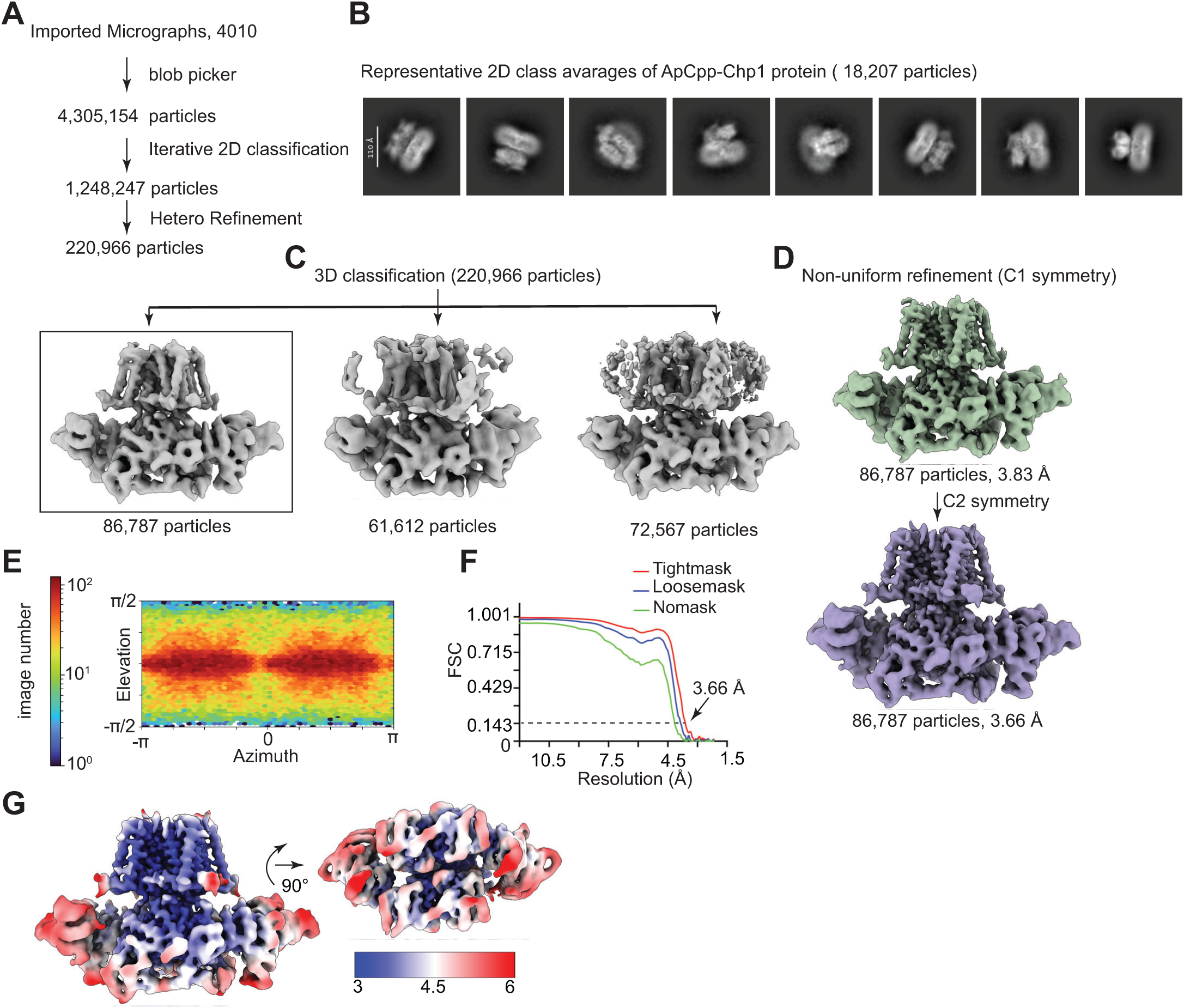
Cryo-EM data processing workflow for ApCpp bound Chp1. **(A)** Schematic summary of particle picking and subsequent selection steps. **(B)** Representative 2D class averages of ApCpp-bound Chp1, showing distinct structural features. **(C)** 3D classification of curated particles, resulting in multiple structural classes; class chosen for further analysis is indicated. **(D)** Final 3D density maps after non-uniform refinement. **(E)** Distribution of particle orientations used in the final reconstruction. **(F)** Fourier shell correlation (FSC) plots from the non-uniform refinement procedure, calculated with no mask, a loose mask, and a tight mask to determine overall resolution. **(G)** Local resolution maps of the refined structures, with resolution values expressed in angstroms (Å).

**Figure S10.**
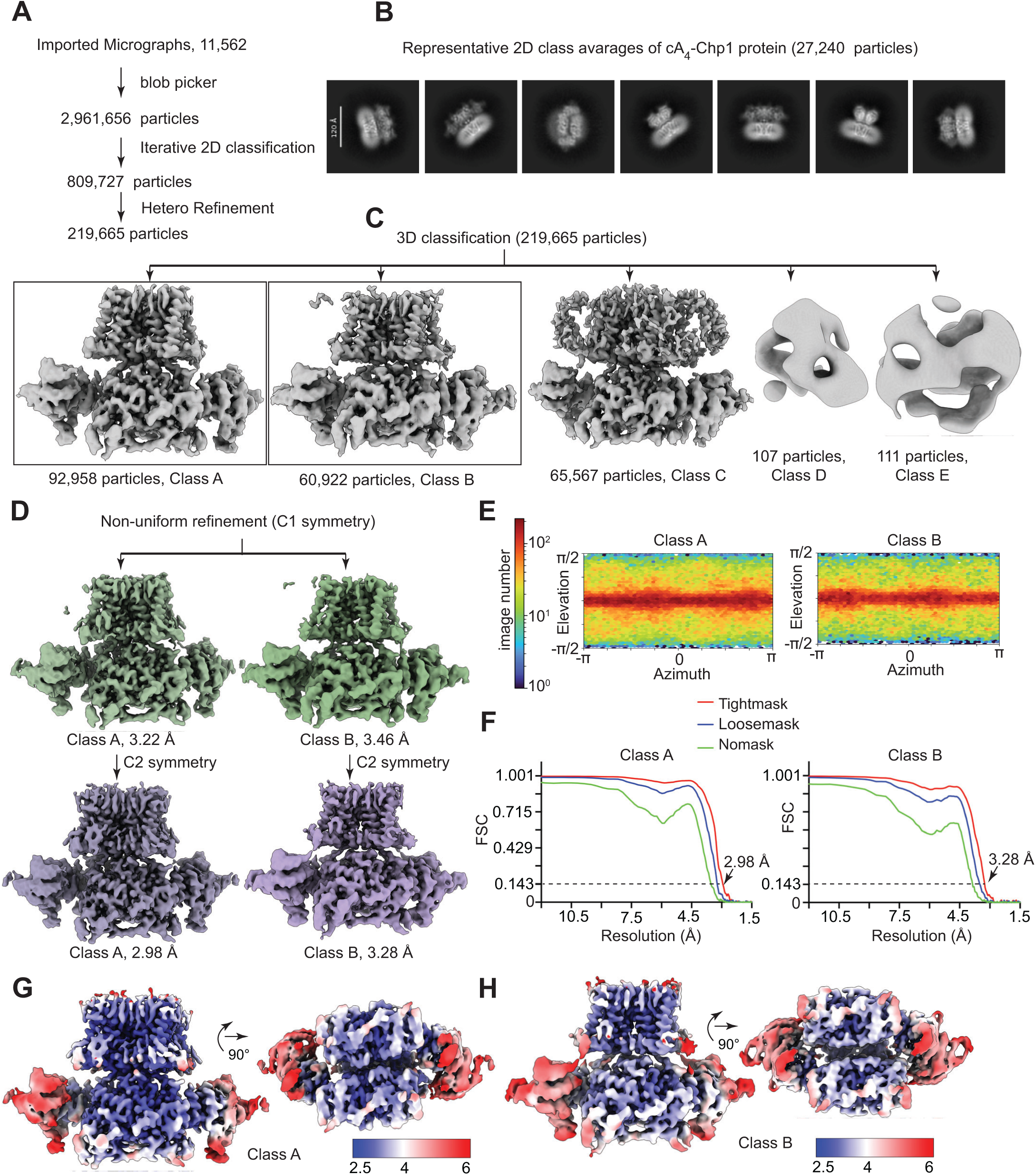
Cryo-EM data processing workflow for cA_4_ bound Chp1. **(A)** Workflow outlining particle picking and selection. **(B)** Representative 2D class averages of cA4-bound Chp1, showing well-resolved structural features. **(C)** 3D classification of selected particles into multiple conformational classes; classes retained for downstream refinement are indicated. **(D)** Final 3D reconstructions obtained following non-uniform refinement. **(E)** Angular distribution of particles contributing to the final maps. **(F)** Fourier shell correlation (FSC) curves derived from the non-uniform refinement job, calculated with no mask, a loose mask, and a tight mask to evaluate global resolution. **(G-H)** Local resolution distributions of the refined density maps, with resolution values reported in angstroms (Å).

**Figure S11.**
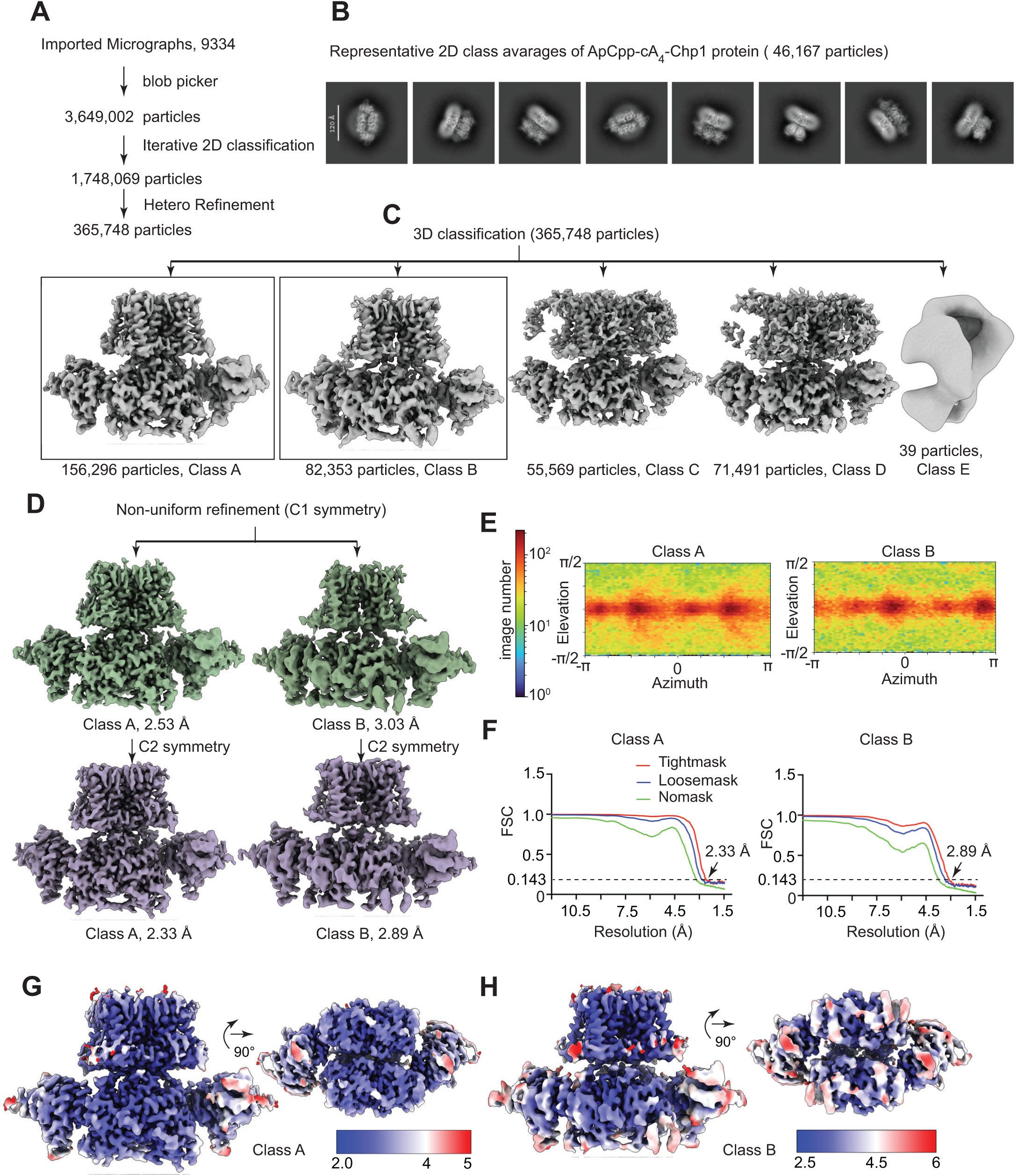
Cryo-EM data processing workflow for cA_4_ and ApCpp bound Chp1. **(A)** Overview of particle picking and subsequent particle cleaning steps. **(B)** Representative 2D class averages of the cA_4_ and ApCpp–bound Chp1 complex. **(C)** 3D classification of selected particles into multiple structural classes; classes chosen for further refinement are indicated. **(D)** Final 3D density maps following non-uniform refinement. **(E)** Angular distribution of particles contributing to the final reconstruction. **(F)** Fourier shell correlation (FSC) curves calculated during non-uniform refinement using no mask, a loose mask, and a tight mask to assess map resolution. **(G-H)** Local resolution maps of the refined structures, with resolution values shown in angstroms (Å).

**Table S1.**
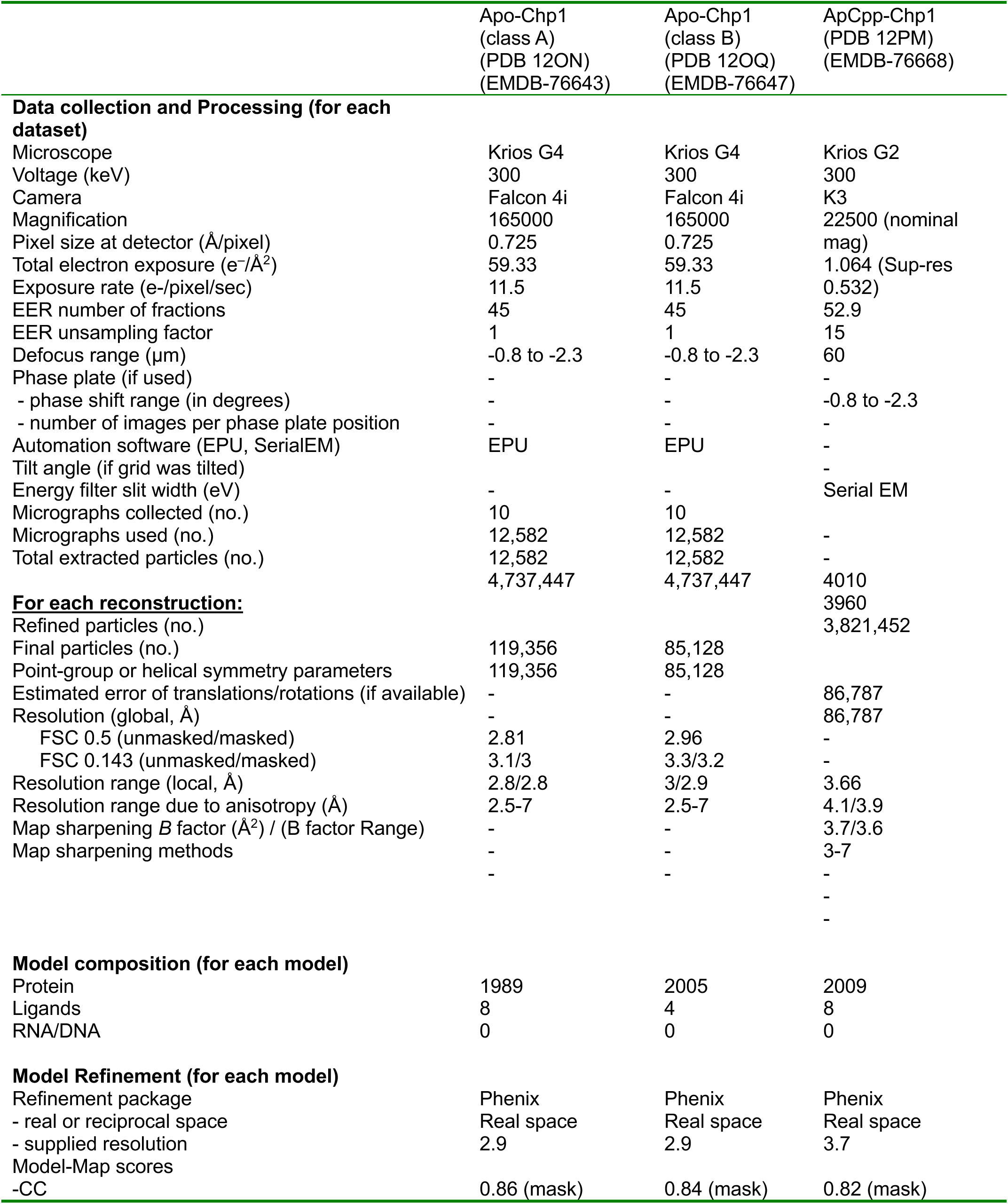
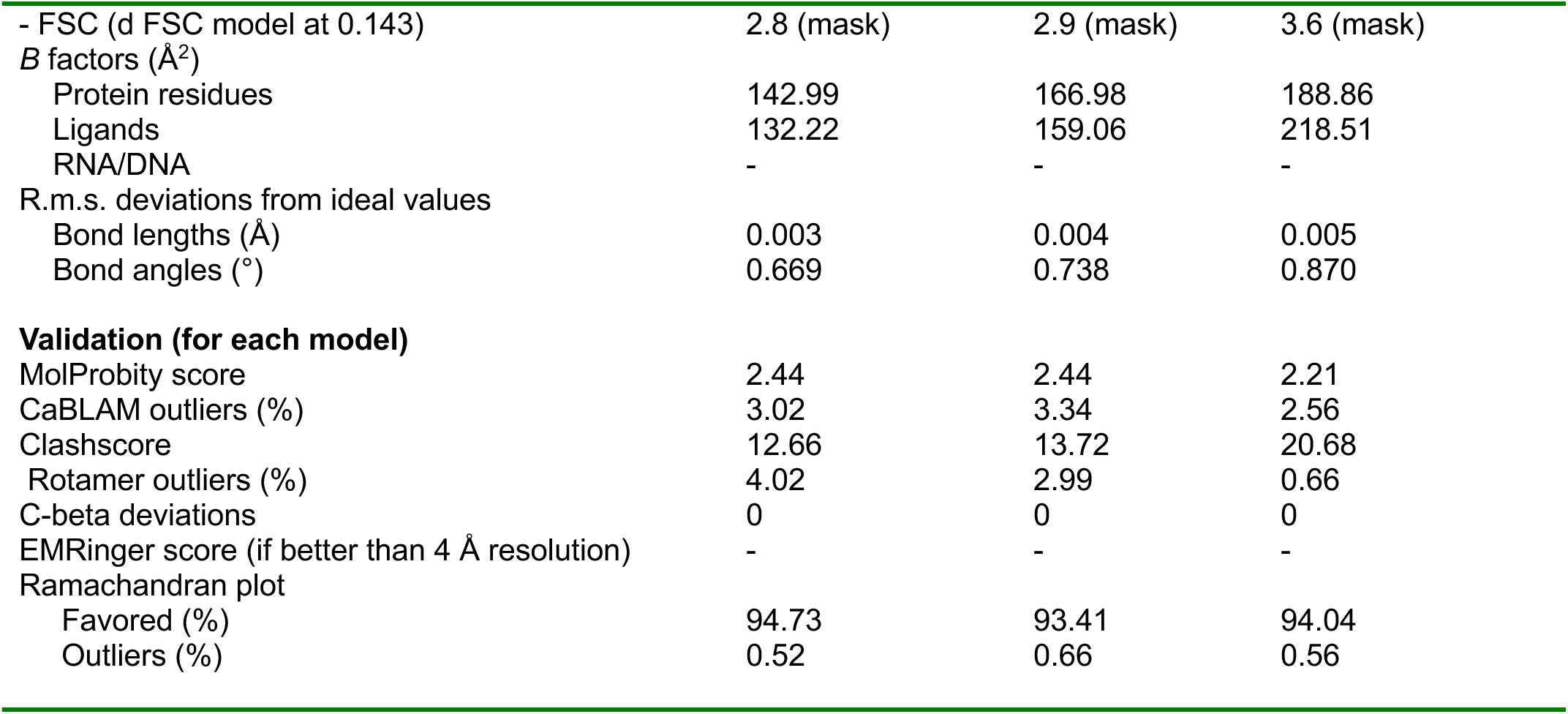
Cryo-EM data collection, refinement, and validation statistics for apo Chp1 and ApCpp-bound Chp1.

**Table S2.**
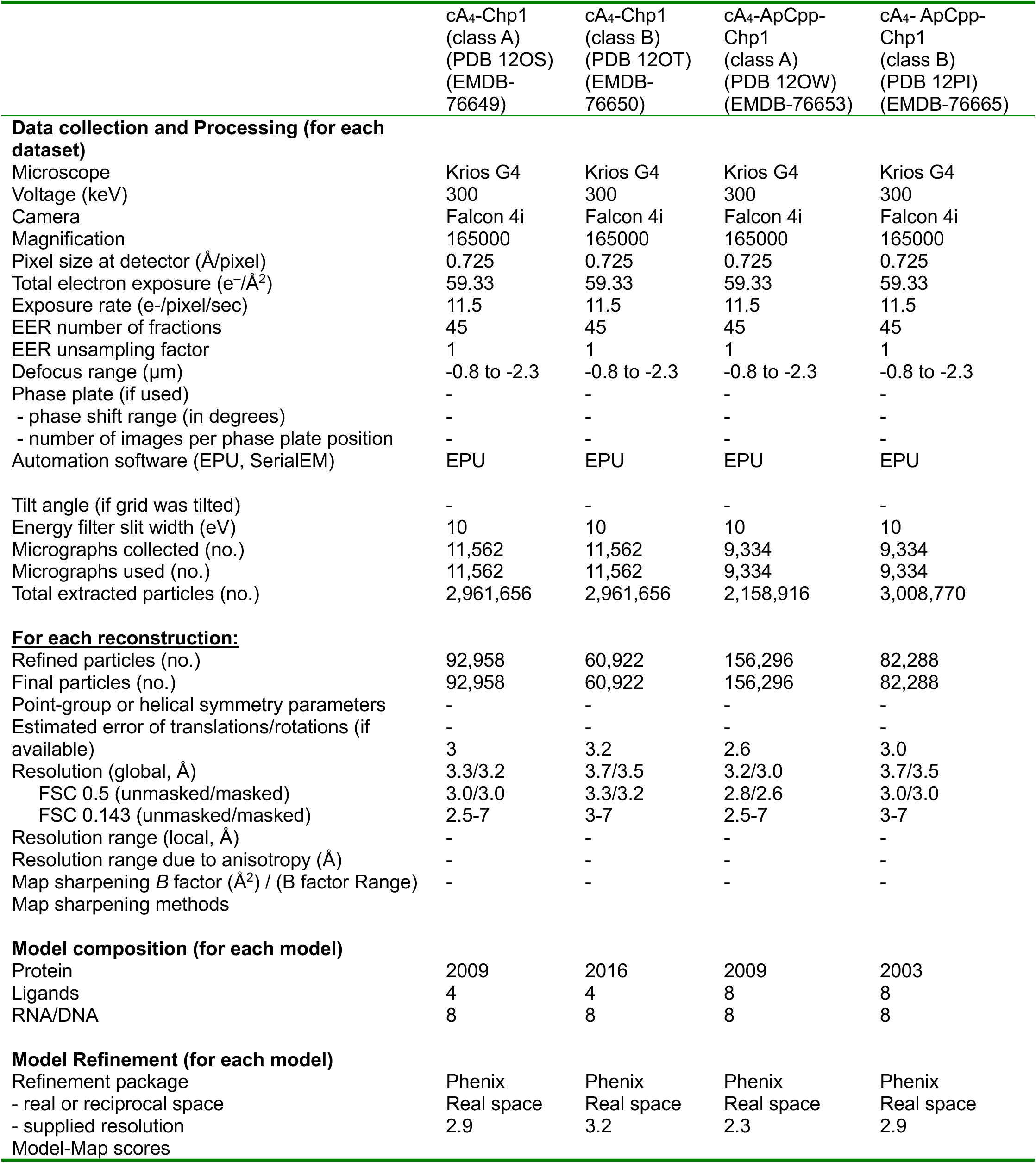
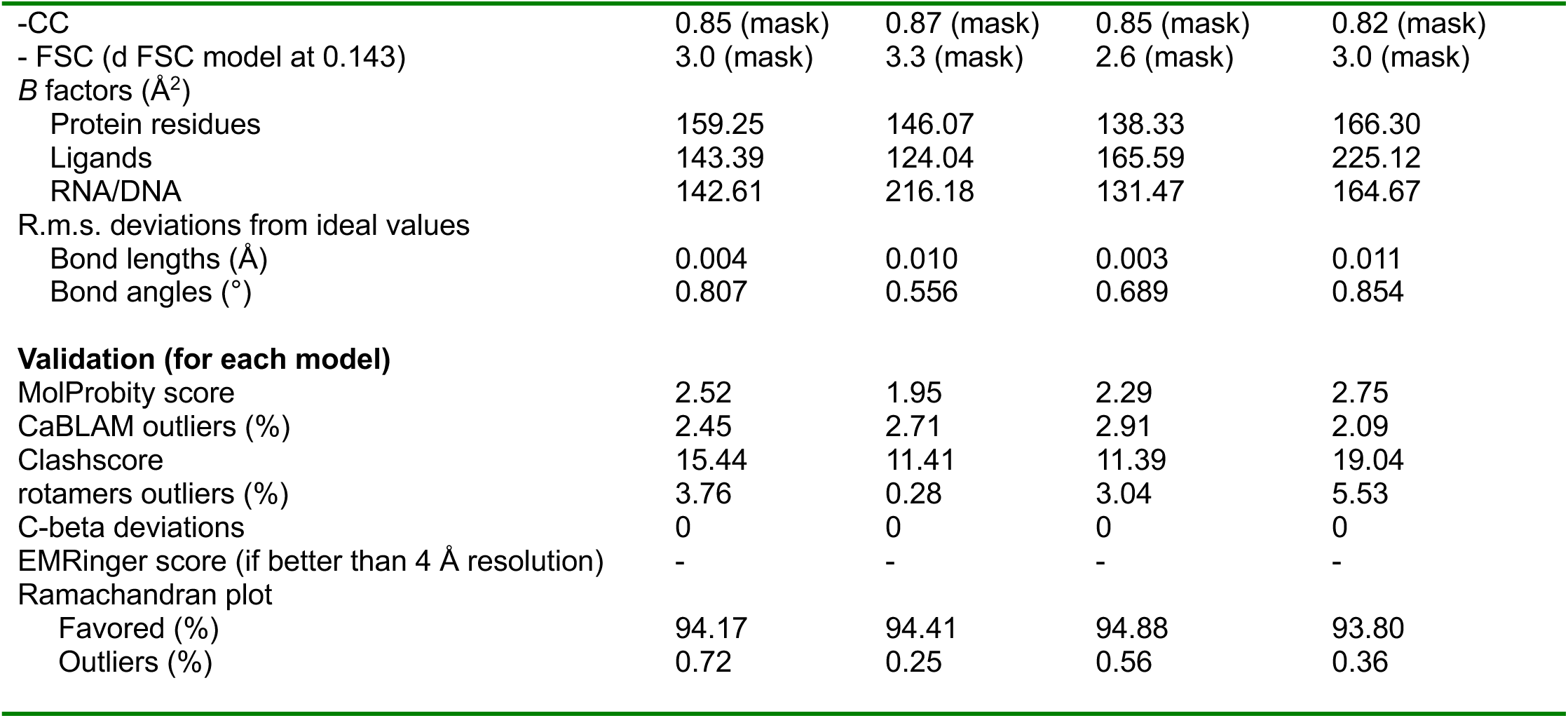
Cryo-EM data collection, refinement, and validation statistics for cA_4_-bound Chp1 and cA_4_-ApCpp-bound Chp1.

## REFERENCES

1. M. Bramkamp, D. Scheffers, Bacterial membrane dynamics: Compartmentalization and repair. Mol. Microbiol. 120, 490–501 (2023).

2. X. Cheng, J. C. Smith, Biological Membrane Organization and Cellular Signaling. Chem. Rev. 119, 5849–5880 (2019).

3. A. L. Davidson, J. Chen, ATP-binding cassette transporters in bacteria. Annu. Rev. Biochem. 73, 241–268 (2004).

4. G. R. Dubyak, Ion homeostasis, channels, and transporters: an update on cellular mechanisms. Adv. Physiol. Educ. 28, 143–154 (2004).

5. B. Martinac, Y. Saimi, C. Kung, Ion Channels in Microbes. Physiol. Rev. 88, 1449–1490 (2008).

6. L. Abdul Kadir, M. Stacey, R. Barrett-Jolley, Emerging Roles of the Membrane Potential: Action Beyond the Action Potential. Front. Physiol. 9, 1661 (2018).

7. J. M. Benarroch, M. Asally, The Microbiologist’s Guide to Membrane Potential Dynamics. Trends Microbiol. 28, 304–314 (2020).

8. B. Duncan-Lowey, Effector-mediated membrane disruption controls cell death in CBASS antiphage defense.

9. C. F. Baca, Y. Yu, J. T. Rostøl, P. Majumder, D. J. Patel, L. A. Marraffini, The CRISPR effector Cam1 mediates membrane depolarization for phage defence. Nature 625, 797–804 (2024).

10. Cap1 forms a cyclic tetra-adenylate-induced membrane pore during the type III-A CRISPR-Cas immune response.

11. N. Tal, B. R. Morehouse, A. Millman, A. Stokar-Avihail, C. Avraham, T. Fedorenko, E. Yirmiya, E. Herbst, A. Brandis, T. Mehlman, Y. Oppenheimer-Shaanan, A. F. A. Keszei, S. Shao, G. Amitai, P. J. Kranzusch, R. Sorek, Cyclic CMP and cyclic UMP mediate bacterial immunity against phages. Cell 184, 5728–5739.e16 (2021).

12. A. Millman, A. Bernheim, A. Stokar-Avihail, T. Fedorenko, M. Voichek, A. Leavitt, Y. Oppenheimer-Shaanan, R. Sorek, Bacterial Retrons Function In Anti-Phage Defense. Cell 183, 1551-1561.e12 (2020).

13. R. Barrangou, C. Fremaux, H. Deveau, M. Richards, P. Boyaval, S. Moineau, D. A. Romero, P. Horvath, CRISPR provides acquired resistance against viruses in prokaryotes. Science 315, 1709–1712 (2007).

14. L. A. Marraffini, E. J. Sontheimer, CRISPR interference limits horizontal gene transfer in staphylococci by targeting DNA. Science 322, 1843–1845 (2008).

15. K. S. Makarova, S. A. Shmakov, Y. I. Wolf, P. Mutz, H. Altae-Tran, C. L. Beisel, S. J. J. Brouns, E. Charpentier, D. Cheng, J. Doudna, D. H. Haft, P. Horvath, S. Moineau, F. J. M. Mojica, P. Pausch, R. Pinilla-Redondo, S. A. Shah, V. Siksnys, M. P. Terns, J. Tordoff, Č. Venclovas, M. F. White, A. F. Yakunin, F. Zhang, R. A. Garrett, R. Backofen, J. van der Oost, R. Barrangou, E. V. Koonin, An updated evolutionary classification of CRISPR-Cas systems including rare variants. Nat. Microbiol. 10, 3346–3361 (2025).

16. G. Stella, L. Marraffini, Type III CRISPR-Cas: beyond the Cas10 effector complex. Trends Biochem. Sci. 49, 28–37 (2024).

17. S. Gruschow, S. McQuarrie, K. Ackermann, S. McMahon, B. E. Bode, T. M. Gloster, M. F. White, CRISPR antiphage defence mediated by the cyclic nucleotide-binding membrane protein Csx23. Nucleic Acids Res 52, 2761–2775 (2024).

18. C. F. Baca, Y. Yu, J. T. Rostol, P. Majumder, D. J. Patel, L. A. Marraffini, The CRISPR effector Cam1 mediates membrane depolarization for phage defence. Nature 625, 797–804 (2024).

19. P. Majumder, C. W. Cahir, C. G. Roberts, D. J. Patel, L. A. Marraffini, Cap1 forms a cyclic tetra-adenylate-induced membrane pore during the type III-A CRISPR-Cas immune response. bioRxiv: The Preprint Server for Biology 2025.11.13.688252 [Preprint] (2025). 10.1101/2025.11.13.688252.

20. O. Niewoehner, C. Garcia-Doval, J. T. Rostøl, C. Berk, F. Schwede, L. Bigler, J. Hall, L. A. Marraffini, M. Jinek, Type III CRISPR–Cas systems produce cyclic oligoadenylate second messengers. Nature 548, 543–548 (2017).

21. M. Kazlauskiene, G. Kostiuk, Č. Venclovas, G. Tamulaitis, V. Siksnys, A cyclic oligonucleotide signaling pathway in type III CRISPR-Cas systems. Science 357, 605–609 (2017).

22. G. Stella, L. Ye, S. F. Brady, L. Marraffini, CARF-HAD phosphatase effectors provide immunity during the type III-A CRISPR–Cas response. Nucleic Acids Res. 53, gkaf1363 (2025).

23. J. T. Rostol, L. A. Marraffini, Non-specific degradation of transcripts promotes plasmid clearance during type III-A CRISPR-Cas immunity. Nat Microbiol 4, 656–662 (2019).

24. J. Payandeh, T. Scheuer, N. Zheng, W. A. Catterall, The crystal structure of a voltage-gated sodium channel. Nature 475, 353–358 (2011).

25. S. G. Brohawn, J. Del Mármol, R. MacKinnon, Crystal Structure of the Human K2P TRAAK, a Lipid- and Mechano-Sensitive K^+^ Ion Channel. Science 335, 436–441 (2012).

26. C. Boiteux, I. Vorobyov, T. W. Allen, Ion conduction and conformational flexibility of a bacterial voltage-gated sodium channel. Proc. Natl. Acad. Sci. 111, 3454–3459 (2014).

27. W. A. Catterall, M. J. Lenaeus, T. M. Gamal El-Din, Structure and Pharmacology of Voltage-Gated Sodium and Calcium Channels. Annu. Rev. Pharmacol. Toxicol. 60, 133–154 (2020).

28. J. Huang, X. Pan, N. Yan, Structural biology and molecular pharmacology of voltage-gated ion channels. Nat. Rev. Mol. Cell Biol. 25, 904–925 (2024).

29. J. M. Jarodsky, J. B. Myers, S. L. Reichow, Reversible lipid-mediated pH-gating of connexin-46/50 by cryo-EM. Nat. Commun. 17, 1606 (2026).

30. T. M. Gamal El-Din, M. J. Lenaeus, N. Zheng, W. A. Catterall, Fenestrations control resting-state block of a voltage-gated sodium channel. Proc. Natl. Acad. Sci. 115, 13111–13116 (2018).

31. L. Coronel, G. Di Muccio, B. S. Rothberg, A. Giacomello, V. Carnevale, Lipid-mediated hydrophobic gating in the BK potassium channel. Nat. Commun. 16, 7354 (2025).

32. M. Lichtenegger, O. Tiapko, B. Svobodova, T. Stockner, T. N. Glasnov, W. Schreibmayer, D. Platzer, G. G. De La Cruz, S. Krenn, R. Schober, N. Shrestha, R. Schindl, C. Romanin, K. Groschner, An optically controlled probe identifies lipid-gating fenestrations within the TRPC3 channel. Nat. Chem. Biol. 14, 396–404 (2018).

33. Y. Y. Dong, A. C. W. Pike, A. Mackenzie, C. McClenaghan, P. Aryal, L. Dong, A. Quigley, M. Grieben, S. Goubin, S. Mukhopadhyay, G. F. Ruda, M. V. Clausen, L. Cao, P. E. Brennan, N. A. Burgess-Brown, M. S. P. Sansom, S. J. Tucker, E. P. Carpenter, K2P channel gating mechanisms revealed by structures of TREK-2 and a complex with Prozac. Science 347, 1256–1259 (2015).

34. S. Grüschow, S. McQuarrie, K. Ackermann, S. McMahon, B. E. Bode, T. M. Gloster, M. F. White, CRISPR antiphage defence mediated by the cyclic nucleotide-binding membrane protein Csx23. Nucleic Acids Res. 52, 2761–2775 (2024).

35. D. Novo, N. G. Perlmutter, R. H. Hunt, H. M. Shapiro, Accurate flow cytometric membrane potential measurement in bacteria using diethyloxacarbocyanine and a ratiometric technique. Cytometry 35, 55–63 (1999).

36. G. W. Goldberg, W. Jiang, D. Bikard, L. A. Marraffini, Conditional tolerance of temperate phages via transcription-dependent CRISPR-Cas targeting. Nature 514, 633–637 (2014).

37. P. Mitchell, Chemiosmotic coupling in oxidative and photosynthetic phosphorylation. Biol. Rev. Camb. Philos. Soc. 41, 445–502 (1966).

38. M. A. Farha, C. P. Verschoor, D. Bowdish, E. D. Brown, Collapsing the proton motive force to identify synergistic combinations against Staphylococcus aureus. Chem. Biol. 20, 1168–1178 (2013).

39. N. Ayoub, A. Gedeon, H. Munier-Lehmann, A journey into the regulatory secrets of the de novo purine nucleotide biosynthesis. Front. Pharmacol. 15, 1329011 (2024).

40. B. N. Kreiswirth, S. Löfdahl, M. J. Betley, M. O’Reilly, P. M. Schlievert, M. S. Bergdoll, R. P. Novick, The toxic shock syndrome exotoxin structural gene is not detectably transmitted by a prophage. Nature 305, 709–712 (1983).

41. A. Punjani, J. L. Rubinstein, D. J. Fleet, M. A. Brubaker, cryoSPARC: algorithms for rapid unsupervised cryo-EM structure determination. Nat. Methods 14, 290–296 (2017).

42. E. F. Pettersen, T. D. Goddard, C. C. Huang, G. S. Couch, D. M. Greenblatt, E. C. Meng, T. E. Ferrin, UCSF Chimera—A visualization system for exploratory research and analysis. J. Comput. Chem. 25, 1605–1612 (2004).

43. P. Emsley, K. Cowtan, *Coot* : model-building tools for molecular graphics. Acta Crystallogr. D Biol. Crystallogr. 60, 2126–2132 (2004).

44. P. V. Afonine, B. K. Poon, R. J. Read, O. V. Sobolev, T. C. Terwilliger, A. Urzhumtsev, P. D. Adams, Real-space refinement in *PHENIX* for cryo-EM and crystallography. Acta Crystallogr. Sect. Struct. Biol. 74, 531–544 (2018).

45. J. Abramson, J. Adler, J. Dunger, R. Evans, T. Green, A. Pritzel, O. Ronneberger, L. Willmore, A. J. Ballard, J. Bambrick, S. W. Bodenstein, D. A. Evans, C.-C. Hung, M. O’Neill, D. Reiman, K. Tunyasuvunakool, Z. Wu, A. Žemgulytė, E. Arvaniti, C. Beattie, O. Bertolli, A. Bridgland, A. Cherepanov, M. Congreve, A. I. Cowen-Rivers, A. Cowie, M. Figurnov, F. B. Fuchs, H. Gladman, R. Jain, Y. A. Khan, C. M. R. Low, K. Perlin, A. Potapenko, P. Savy, S. Singh, A. Stecula, A. Thillaisundaram, C. Tong, S. Yakneen, E. D. Zhong, M. Zielinski, A. Žídek, V. Bapst, P. Kohli, M. Jaderberg, D. Hassabis, J. M. Jumper, Accurate structure prediction of biomolecular interactions with AlphaFold 3. Nature 630, 493–500 (2024).

